# Aberrant homeodomain-DNA cooperative dimerization underlies distinct developmental defects in two dominant *CRX* retinopathy models

**DOI:** 10.1101/2024.03.12.584677

**Authors:** Yiqiao Zheng, Gary D. Stormo, Shiming Chen

**Affiliations:** Molecular Genetics and Genomics Graduate Program, Division of Biological and Biomedical Sciences, Washington University in St Louis, Saint Louis, Missouri, 63110, USA; Department of Ophthalmology and Visual Sciences, Washington University in St Louis, Saint Louis, Missouri, 63110, USA; Department of Developmental Biology, Washington University in St Louis, Saint Louis, Missouri, 63110, USA; Department of Genetics, Washington University in St Louis, Saint Louis, Missouri, 63110, USA

**Keywords:** CRX, homeodomain, transcription factor, missense mutations, DNA binding specificity, DNA binding cooperativity, inherited retinal disease, photoreceptor development, chromatin remodeling

## Abstract

Paired-class homeodomain transcription factors (HD TFs) play essential roles in vertebrate development, and their mutations are linked to human diseases. One unique feature of paired-class HD is cooperative dimerization on specific palindrome DNA sequences. Yet, the functional significance of HD cooperative dimerization in animal development and its dysregulation in diseases remain elusive. Using the retinal TF Cone-rod Homeobox (CRX) as a model, we have studied how blindness-causing mutations in the paired HD, p.E80A and p.K88N, alter CRX’s cooperative dimerization, lead to gene misexpression and photoreceptor developmental deficits in dominant manners. CRX^E80A^ maintains binding at monomeric WT CRX motifs but is deficient in cooperative binding at dimeric motifs. CRX^E80A^’s cooperativity defect impacts the exponential increase of photoreceptor gene expression in terminal differentiation and produces immature, non-functional photoreceptors in the *Crx*^*E80A*^ retinas. CRX^K88N^ is highly cooperative and localizes to ectopic genomic sites with strong enrichment of dimeric HD motifs. CRX^K88N^’s altered biochemical properties disrupt CRX’s ability to direct dynamic chromatin remodeling during development to activate photoreceptor differentiation programs and silence progenitor programs. Our study here provides *in vitro* and *in vivo* molecular evidence that paired-class HD cooperative dimerization regulates neuronal development and dysregulation of cooperative binding contributes to severe dominant blinding retinopathies.

## Introduction

Homeodomain transcription factors (HD TFs) are essential for diverse biological processes in vertebrate development, including body plan specification, pattern formation, and cell fate specification (1-3). Paradoxically, for a protein domain that has evolved numerous functional specificities, it binds with high affinity to closely related DNA motifs that are typically only 5-6 base pairs long (4-7). It has thus been proposed that additional mechanisms are required to achieve the individual functions of homeoproteins (8).

The paired-class HD family possesses a unique feature in that members of this class confer cooperative dimerization on specific dimeric DNA sequences (8). The “cooperative” interaction here is where the first HD-DNA half-site binding event greatly enhances the binding of a second molecule to the other half-site (Figure 1A). The paired-class HD cooperative dimerization solely relies on the 60-amino-acid HD, distinguishing it from the other HD families that require additional domains to form higher-order DNA binding complexes (8,9). In addition, the recognition helix residues that determine paired-class HD’s DNA binding specificity at high-affinity monomeric motifs also confer distinct cooperative dimerization properties, including the preferred spacer length and identity between the two 5’-TAAT-3’ core motifs and the magnitude of cooperativity (8,10). Yet, the functional importance of paired-class HD cooperative dimerization in development and its dysregulation in human diseases remains elusive.

**Figure 1.**
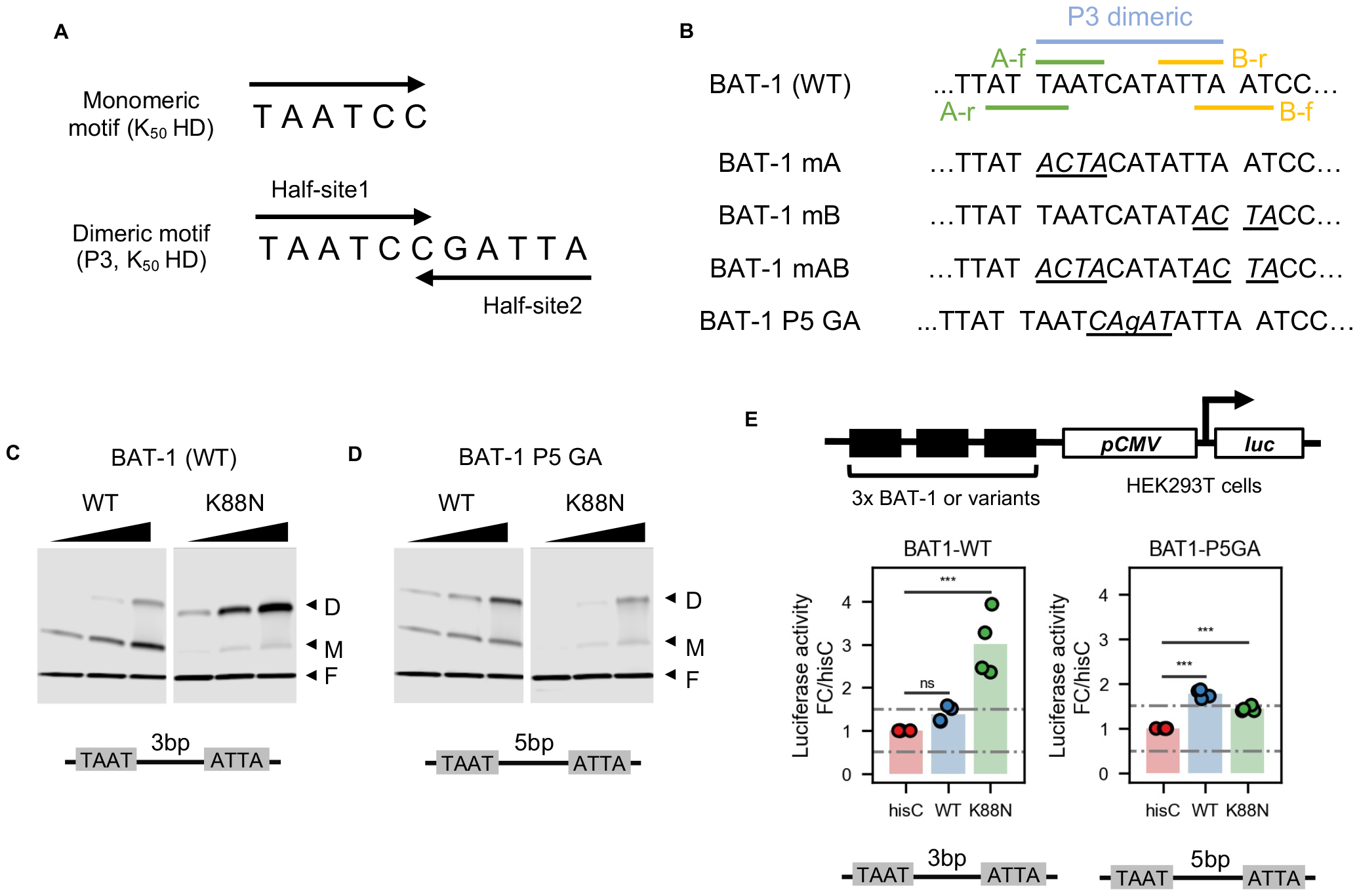
K88N mutation significantly increased CRX HD’s cooperative binding and transactivation activity on BAT-1 sequence containing a P3 dimeric HD motif. **A**. Diagrams depicting K_50_ HD preferred monomeric and dimeric P3 motifs. **B**. Alignments showing WT BAT-1 probe sequences and variants. The P3 dimeric HD motif and four monomeric HD core motifs 5’-TAAT-3’ are labelled. f and r indicates whether the core motif is on the forward or reverse strand. In BAT-1 variants, the mutated nucleotides are italicized and underlined. **C.&D**. EMSA gel images showing increasing amount of WT or K88N HD peptides bound to a fixed amount of BAT-1 (WT) or P5 GA control probes. The cartoon underneath each gel image shows the dimeric HD motif configuration and is labelled with the spacer length. **E**. Schematics and barcharts of luciferase reporter assays comparing CRX WT and K88N transactivation activity at BAT-1 and variant enhancer sequences. *p-values* of one-way ANOVA are annotated. ns: >5e-2; ***: <=1e-3.

The retina is an excellent model system for dissecting the role of HD TFs in central nervous system (CNS) development and neurological disorders. HD TFs are expressed at various stages of retinal development to pattern the neuroepithelium, specify the fate of specific cell types from multi-potent retinal progenitor cells, and direct differentiation of individual cell types (11-13). Notably, the retina shares general neuronal and anatomical features with the brain, and the molecular pathways uncovered in the retina often inform the programs governing brain development. The accessibility and rich cellular and molecular toolbox make the retina a valuable tool for deciphering HD TF mechanisms of action in development and studying mutations associated with neurological disorders (14).

We have studied CRX, a paired-class HD TF essential for photoreceptor cells in the retina, as a model to understand HD-DNA interactions in normal development and dominant blinding diseases (15-18). Photoreceptors are specialized neurons in the retina that sense light and initiate vision through the phototransduction process. In vertebrates, photoreceptors come in two major classes, rods and cones, that mediate vision in dim and bright light, respectively. Animal studies have demonstrated that *Crx* expression is activated in post-mitotic photoreceptor precursors and maintained throughout adult life (19-21). The disruption of CRX expression or function profoundly impacts photoreceptor development, leading to rapid degeneration of immature, non-functional photoreceptors (18,22,23). Coding sequence mutations in human *CRX* have been associated with at least three inherited retinal diseases (IRDs) that primarily affect photoreceptors: Leber congenital amaurosis 7 (LCA7, OMIM: 613829), Cone-rod dystrophy 2 (CoRD2, OMIM: 120970), retinitis pigmentosa (RP, OMIM: 268000). *CRX*-associated retinopathies vary significantly in the ages of onset, severity, and disease progression (15,16). The phenotype heterogeneity suggests that individual mutation may cause disease via distinct pathogenic mechanisms. Deciphering these mechanisms is, therefore, critical for informing the future development of therapeutic approaches.

CRX has two functional domains – the N-terminal DNA-binding domain (homeodomain, HD) and the C-terminal transcription effector domain; both are required for target gene regulation in photoreceptor development and maintenance. Disease-causing mutations are distributed across both functional domains, with amino acid substitutions primarily observed in the CRX HD (15,16). To understand how HD mutations alter CRX-DNA interactions in development and lead to photoreceptor diseases, we have previously reported two human mutation *knock-in* mouse models (17), each carrying a gain-of-function mutation, p.E80A (E80A) and p.K88N (K88N), that are associated with dominant LCA and dominant CoRD, respectively. Using an integrated approach that combines quantitative *in vitro* biochemical models, functional genomics, cellular profiling, and functional testing in mouse models, we found that E80A and K88N alter CRX DNA binding specificity and lead to distinct photoreceptor deficits in mutant mouse retinas. Yet, the proposed mechanisms were primarily based on analyzing CRX HD-DNA interactions at monomeric HD motifs both *in vitro* and *in vivo*. Given that CRX is a paired-class HD TF, it is unclear whether E80A and K88N mutations affect CRX HD cooperative dimerization and how mutant CRX activity interferes with WT CRX functions when both alleles are present, leading to the severe dominant photoreceptor deficits in developing mouse retinas.

Here, we extend our multi-omics approach to further elucidate the consequences of E80A and K88N mutations on CRX regulatory activities at both monomeric and dimeric HD DNA motifs and on the photoreceptor developmental programs. Using Coop-seq, a high-throughput *in vitro* HD-DNA binding cooperativity assay, we compared the profile of each mutant protein to that of WT CRX and discovered unique cooperativity changes. We performed ATAC-seq analysis of WT, heterozygous, and homozygous *Crx* mutant mouse retinas and identified shared and genotype-specific chromatin landscape changes. Motif analysis found enrichment of distinct monomeric and dimeric HD DNA motifs in differentially accessible regions of each mutant model, consistent with the specific DNA binding cooperativity changes observed *in vitro*. Importantly, integrated analysis of the epigenome and transcriptome data revealed a correlation between changes in local chromatin accessibility and gene mis-expression, explaining the genotype-specific photoreceptor developmental deficits *in vivo*. Our study affords insights into the distinct pathogenic mechanisms of dominant CRX HD mutations and emphasizes the importance of precise HD-DNA interactions in stage-specific transcriptional regulation during photoreceptor development.

## Results

### CRX K88N but not WT or E80A HD confers strong cooperative dimerization on p*Rho BAT-1* probe

As a paired-class HD TF, CRX binds 6mer monomeric DNA motifs with high affinity (Figure 1A top) and demonstrates cooperative binding to specific dimeric HD motifs, historically known as P3 sequences (8,10) (Figure 1A bottom). In a P3 sequence, the two half-site core motifs 5’-TAAT-3’ are separated by a 3bp (base pair) spacer and form an approximate palindrome. The length and identity of the spacer are determined by amino acids in the homeodomain recognition helix (CRX aa.80-96). Distinctively, in the P3 configuration, the two 6mer monomeric half-sites overlap by a single base pair, which has been predicted to elicit the cooperative mode of binding. In the previous report, we found that disease-causing mutations E80A, K88N, and R90W, all located within the CRX HD recognition helix, differentially affect CRX HD DNA-binding specificity at 6mer monomeric sequences (17). Here, we asked whether any of the three mutations affect CRX HD cooperative binding at dimeric P3 sequences using electrophoretic mobility shift assays (EMSAs).

We chose the *BAT-1* fragment as an EMSA probe since it is an established model template to assay CRX HD’s dimeric DNA binding (20,26) (Figure 1B). The *BAT-1* sequence is a fragment in the promoter of *rhodopsin*, a gene that encodes the rod-specific photopigment and is a direct target of CRX *in vivo*. The *BAT-1* fragment contains a central dimeric P3 sequence TAAT<u>CAT</u>ATTA and additional overlapping monomeric HD motifs (Figure 1B). We also generated a series of *BAT-1* variant fragments to explicitly interrogate the interactions between the two half-sites (Figure 1B). Specifically, in the *BAT-1* P5 GA variant, the two half-sites constituting the P3 dimeric sequence are separated by a guanine (G) nucleotide, with each 6mer sequence preserved. The P5 sequence configuration (5bp spacer) has been shown to abolish cooperative dimerization of paired-class homeoproteins (8) and thus was used as a control to visualize non-cooperative/additive dimeric binding events.

WT HD bound strongly to *BAT-1* (WT) and P5 probes as monomeric and dimeric complexes but demonstrated no obvious cooperativity (Figures 1C-1D), as exemplified by the saturation of the monomeric band (M) before the gradual formation of the dimeric band (D). The difference in WT HD bound monomeric *versus* dimeric band intensities between P3 and P5 probes may be attributed to DNA shape features of the two half-sites that intrinsically affect HD-DNA interactions without explicitly changing the underlying sequences (32,33). Surprisingly, K88N HD showed strong dimeric binding with weak monomeric binding at the *BAT-1* (WT) probe (Figures 1C-1D, S1A-S1B), characteristic of cooperative dimerization where the binding of one molecule stimulates the binding of a second molecule (8,10). Increasing the spacer length (*BAT-1* P5 GA, Figure 1D) or abolishing either half-site of the P3 sequence (*BAT-1* mA and mB, Figure S1C) resulted in diminished K88N HD dimeric DNA binding, corroborating the essential P3 configuration and intact palindromic half-sites in mediating K88N HD cooperative dimerization. The diminished K88N HD dimeric binding at *BAT-1* P5, mA, and mB probes unequivocally argues that K88N mutation does not render CRX K88N proteins obligatory dimers, and cooperative dimerization is mediated through specific HD-DNA interactions. In comparison, E80A HD bound stronger as monomeric complexes and weaker as dimeric complexes at both *BAT-1* (WT) and P5 probes when compared to WT HD (Figures S1A-S1B), which may inherently relate to E80A HD’s reduced binding specificity at monomeric sequences (17); R90W HD bound poorly to either *BAT-1* (WT) or P5 probe, consistent with R90W being a loss-of-function mutation that reduces CRX HD’s overall DNA binding affinity (26). In summary, only CRX K88N but not WT or E80A HD confers strong cooperative dimerization at the *BAT-1* probe containing a P3 dimeric HD motif.

### CRX K88N cooperative dimerization mediates strong reporter gene activation in HEK293T cells

To determine the consequences of K88N HD-gained cooperative dimerization on CRX’s transactivation activity, we performed luciferase reporter assays in the HEK293T cells with enhancers harboring three tandem repeats of *BAT-1* or variant sequences identical to that used in EMSAs. Consistent with EMSA results, CRX K88N demonstrated strong activator activity at 3x*BAT-1* WT enhancer harboring the intact P3 dimeric sequence but much weaker activity at all *BAT-1* variants (Figures 1E, S1D). Thus, CRX K88N is a competent transcription activator and CRX K88N’s cooperative dimerization on *BAT-1* P3 sequence mediates strong gene activation likely through stabilizing the dimeric binding complexes.

Interestingly, we found that WT CRX only weakly activated the 3x*BAT-1* WT and mB enhancers and contrastingly activated 3x*BAT-1* P5 and mA enhancers strongly (Figures 1E, S1D). Specifically in the *BAT-1* mA sequence, the sub-optimal 5’-TAATCA-3’(A-f) is destroyed, and a single WT CRX consensus monomeric motif 5’-TAATCC-3’ (B-f) remains intact. The strongest transactivation activity from the 3x*BAT*-1 mA enhancer suggests that CRX monomeric binding at the consensus sequence mediates robust gene activation, consistent with HDs’ known high-affinity binding at monomeric motifs. Although WT CRX did not show apparent cooperative dimerization at the *BAT-1* (WT) probe (Figures 1C-1D), it does not rule out WT CRX’s ability to form cooperative dimers on other P3 sequences in the genome, which was further investigated using unbiased and more quantitative measures below.

### K88N enhances but E80A impairs CRX HD’s cooperative dimerization at various P3 sequences *in vitro*

To characterize CRX HD’s DNA binding cooperativity unbiasedly, we adapted a high-throughput *in vitro* assay, Coop-seq, that determines protein-DNA-binding cooperativity by sequencing (34,35) (Figures 2A-2B). Coop-seq relates to our previously employed Spec-seq in that they both are based on the traditional EMSAs, which allows the physical separation of distinct binding complexes. Coop-seq enables us to accurately measure the cooperativity parameters – interactions between two HD half-sites – for a library of dimeric HD DNA motifs in parallel and quantitatively compare the cooperativity of different CRX HDs. Based on HD-DNA interaction models and previous Spec-seq-generated monomeric CRX HD-DNA binding models (17), we designed a Coop-seq library containing all possible dimeric P3 spacer sequences TAAT<u>NNN</u>ATTA (Figure 2C).

**Figure 2.**
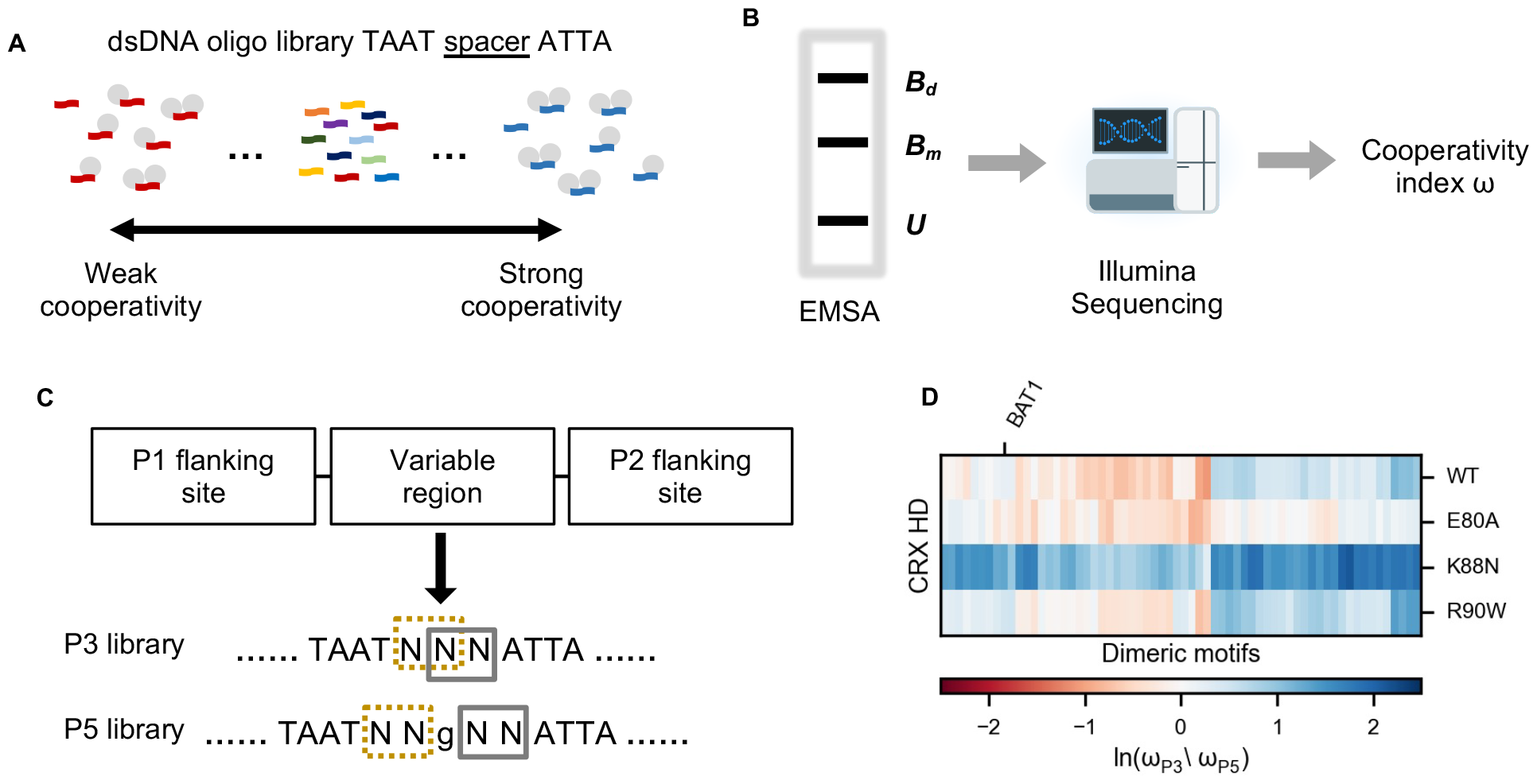
E80A and K88N mutations differently affect CRX HD DNA binding cooperativity at P3 sequences. **A.&B**. Schematics showing the Coop-seq experimental pipelines. dsDNA oligo pools of P3 and/or P5 Coop-seq library are incubated with different HD peptides. The dimeric and monomeric binding complexes are separated from unbound DNAs by EMSA. DNAs are extracted from all three DNA bands and subjected to quantification by Illumina sequencing. B_d_: dimeric band; B_m_: monomeric band; U: unbound band. **C**. Diagram depicting the Coop-seq library design and strategy to match a P3 sequence with a P5 counterpart. Exact oligo sequences can be found in Supplementary Table S1. **D**. Heatmap comparing the relative cooperativity of WT and variant CRX HDs on P3 and P5 libraries (ω_p3_/ω_p5_). Note the relative cooperativity is presented in the Logarithmic scale and ordered by unsupervised hierarchical clustering. The ordered relative cooperativity matrix can be found in Supplementary Table S3.

As a control, we first tested a P5 library (TAAT<u>NN</u>G<u>NN</u>ATTA) (Figure 2C), which is expected to limit cooperativity between half-sites (8,10) (Figures 1D, S1B). We obtained WT and mutant HDs’ DNA binding cooperativity index ω at the P5 library using bacterially-expressed and affinity-purified HD peptides as previously described (Methods). As expected, WT and mutant CRX HDs showed weak cooperativity at P5 sequences (Figures S2A-S2D, Supplementary Table S2). It supports the idea that CRX HDs primarily bind non-overlapping half-sites independently, consistent with homeoproteins’ high affinity binding at monomeric HD motifs.

Next, we compared CRX HDs’ cooperativity profiles on the preferred P3 configuration sequences. To visualize the specific cooperative interactions elicited by the P3 configuration, we normalized the cooperativity index ω at P3 sequences (ω_p3_) by their P5 counterparts (ω_p5_) (Methods, Figure 2C). WT HD showed moderate cooperativity at a small subset of P3 sequences (Figure 2D). Close examination of the P3 sequence spacers of this subset revealed an enrichment of guanine (G) and cytosine (C) bases (Supplementary Table S3), which are known to be preferred by Lys50 (K_50_) HD subfamily of paired-class homeoproteins, including CRX. R90W HD showed a similar cooperativity profile as WT HD with slightly increased cooperativity at a few P3 sequences. This observation is consistent with R90W being a loss-of-function mutation that reduces the overall HD-DNA binding affinity without selectively altering the specific HD-DNA contacts (17,26). E80A HD unexpectedly exhibited significantly reduced cooperativity at nearly all P3 sequences compared to WT HD. Distinctively, K88N HD showed strong DNA-binding cooperativity at all P3 sequences. A previous study found that K_50_ HDs only bind the K_50_ consensus (highest affinity, not necessarily strongest cooperativity) P3 sequence with a cooperativity index of 20-30 (8). In comparison, Glu50 (Q_50_) HDs bind the Q_50_ consensus P3 sequence with a cooperativity index of 240-300. Thus, the enhanced K88N HD cooperativity may relate to its binding specificity change that mimics a Q_50_ HD, as shown in our previous study (17).

Collectively, *in vitro* protein-DNA binding results indicate that E80A mutation impairs CRX HD’s DNA binding cooperativity at specific dimeric HD motifs while K88N mutation drastically alters both the specificity and cooperativity. The luciferase reporter assays highlight the functional distinctions between cooperative dimeric binding at P3 sequences in contrast to non-cooperative dimeric binding at P5 sequences with non-overlapping monomeric half-sites. The specific P3 configuration may have important regulatory implications in development and diseases, which we address using our established animal models below (17).

### *Crx*^*K88N/+*^ and *Crx*^*K88N/N*^ retinas both show severely decreased accessibility at CREs enriched for K_50_ HD motifs

Previous study has established that WT CRX is required to facilitate chromatin remodeling at photoreceptor cis-regulatory elements (CREs) and subsequently activate photoreceptor genes in post-natal retinal development (36). To decipher the roles of CRX’s monomeric binding and cooperative dimeric binding in regulating the photoreceptor epigenome, we performed retinal ATAC-seq on post-natal day 14 (P14) WT, heterozygous and homozygous *Crx* mutant mouse retinas (For concision, we use *Crx*^*E80A*^ and *Crx*^*K88N*^ when both heterozygous and homozygous mutants are being discussed.). We analyzed the ATAC-seq results in conjunction with our previously published P14 CRX ChIP-seq data (17). We asked how E80A and K88N mutation specific changes in HD-DNA interactions affect the chromatin landscape in individual mutant model and how perturbed epigenome relates to photoreceptor gene mis-expression in these retinas.

Our previous study found that CRX WT and K88N prefer different monomeric HD motifs *in vitro* and *in vivo* (17). We thus initially predicted that the *Crx*^*K88N/N*^ retinas show similar chromatin remodeling defects as the loss-of-function *Crx*^*R90W/W*^ model. We also predicted that WT CRX proteins in the *Crx*^*K88N/+*^ retinas can bind canonical CRX binding sites and facilitate chromatin remodeling. Unexpectedly, we found that the *Crx*^*K88N/N*^ retinas showed more significant chromatin accessibility loss at canonical CRX binding sites than the loss-of-function *Crx*^*R90W/W*^ retinas, and the *Crx*^*K88N/+*^ retinas showed reduced accessibility similar to the *Crx*^*R90W/W*^ retinas (Figure 3A). Genomic regions that showed defective chromatin remodeling in the *Crx*^*K88N/+*^ and *Crx*^*K88N/N*^ retinas were enriched for the K_50_ HD motifs (Figure 3B), gain accessibility in normal post-natal retinal development (Figure 3C), and regulate genes important for photoreceptor structures, functions, and maintenance (Figure 3D). The impaired chromatin remodeling at canonical CRX binding sites led to significant photoreceptor gene down-regulation in the developing *Crx*^*K88N/+*^ and *Crx*^*K88N/N*^ retinas (Figure 3E). Thus, defects in chromatin remodeling at regions with K_50_ HD motifs underlie defective photoreceptor differentiation and vision loss in young adults in the *Crx*^*K88N/+*^ and *Crx*^*K88N/N*^ mice (Figure 3F) (17). Since the *Crx*^*R90W/+*^ retinas, expressing a single dose of WT CRX protein, show largely WT phenotypes (18), the severe chromatin remodeling defects and photoreceptor gene mis-expression in the *Crx*^*K88N*^ retinas likely attribute to dominant effects of CRX K88N ectopic activities.

**Figure 3.**
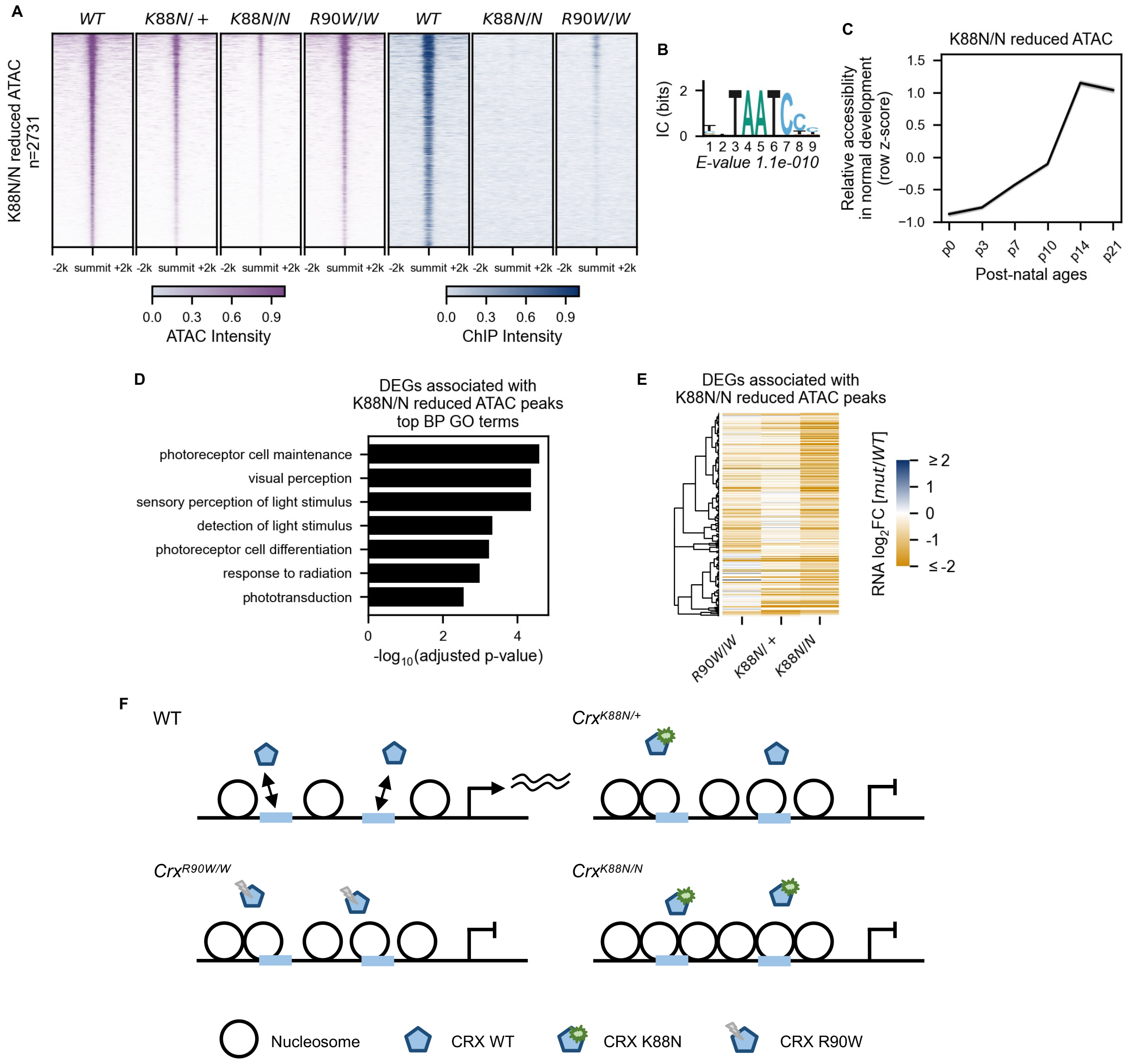
*Crx*^*K88N/+*^ retinas show defective chromatin remodeling at photoreceptor CREs enriched with K_50_ HD motifs. **A**. Heatmaps depicting the normalized ATAC-seq or CRX ChIP-seq signal intensities at *Crx*^*K88N*^-reduced accessible ATAC-seq peaks. **B**. PWM logo and enrichment significance *E-value* of the STREME *de novo* discovered HD motif. **C**. Line plot showing the average developmental accessibility kinetics of *Crx*^*K88N*^-reduced ATAC-seq peaks. The developmental ATAC-seq data is from Aldiri et al. 2017. **D**. Barchart showing Biological Process (BP) Gene Ontology (GO) term enrichment of differentially expressed genes adjacent to *Crx*^*K88N*^-reduced ATAC-seq peaks. **E**. Heatmap comparing the P10 expression changes of *Crx*^*K88N*^-reduced ATAC-seq peaks associated genes in different *Crx* mutant retinas. The gene set is identical to that in **D. F**. Schematics depicting chromatin remodelling defects at photoreceptor regulatory regions in the *Crx*^*K88N/+*^, *Crx*^*K88N/N*^, and *Crx*^*R90W/W*^ retinas.

### *Crx*^*K88N/+*^ and *Crx*^*K88N/N*^ retinas show increased accessibility at CREs enriched for Q_50_ HD motifs

We then sought to understand how CRX K88N ectopic transcription factor activities lead to more severe photoreceptor developmental defects in the *Crx*^*K88N/+*^ and *Crx*^*K88N/N*^ retinas than the loss-of-function *Crx*^*R90W/+*^ and *Crx*^*R90W/W*^ retinas. Different from E80A and R90W which are predicted to affect the stability of HD-DNA binding complex, K88N mutation falls exactly on the DNA binding specificity determining residue of paired-class homeodomains (37-39). Consequently, K88N mutation alters CRX DNA binding preference from that of K_50_ HDs to that resembling Q_50_ HDs (17). *de novo* motif searching on sequences under *Crx*^*K88N*^-increased ATAC-seq peaks (Figure 4A) revealed both dimeric and monomeric Q_50_ HD motifs (Figure 4B). The *Crx*^*K88N*^-increased accessibility ATAC-seq peaks coincided with ectopic CRX K88N binding but showed comparable accessibility in the *Crx*^*R90W/W*^ and WT retinas (Figure 4A). Notably, the *Crx*^*K88N*^ retina ectopically enriched dimeric HD motifs exhibit no strong base preferences within the 3bp spacer (Figure 4B, positions 5-7), which is in line with K88N HD’s strong cooperativity at nearly all P3 Coop-seq library sequences (Figure 2D). Collectively, these observations suggest that ectopic CRX K88N activity at Q_50_ HD motifs rather than loss of WT CRX activity mediates chromatin accessibility gain at ectopic sites in the *Crx*^*K88N/+*^ and *Crx*^*K88N/N*^ retinas that likely contribute to the severe dominant photoreceptor deficits.

**Figure 4.**
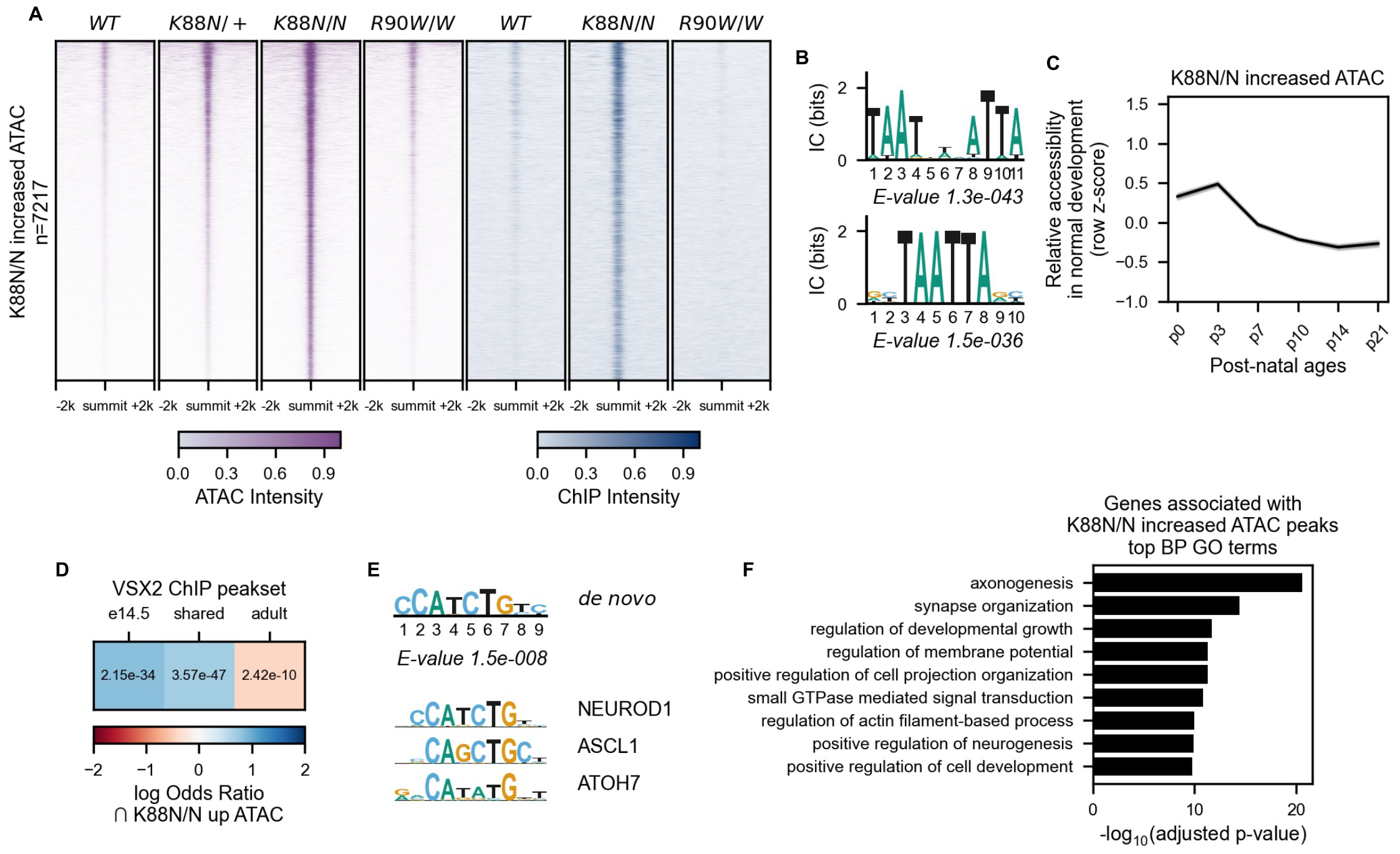
CRX K88N ectopic activity at Q_50_ HD motifs impedes the silencing of progenitor regulatory programs in developing photoreceptors. **A**. Heatmaps depicting the normalized ATAC-seq or CRX ChIP-seq signal intensities at *Crx*^*K88N*^-increased accessible ATAC-seq peaks. **B**. PWM logo and enrichment significance *E-value* of STREME *de novo* discovered HD motifs. **C**. Line plot showing the average developmental accessibility kinetics of *Crx*^*K88N*^-increased ATAC-seq peaks. The developmental ATAC-seq data is from Aldiri et al. 2017. **D**. Heatmap depicting the log odds ratio enrichment of embryonic day 14.5 (e14.5) or adult retinal VSX2 binding sites under *Crx*^*K88N*^-increased ATAC-seq peaks. *p-values* of Fisher’s exact tests are indicated. The VSX2 ChIP-seq data is from Bian et al. 2022. **E**. PWM logo and significance *E-value* of STREME *de novo* discovered basic helix-loop-helix (bHLH) motif under *Crx*^*K88N*^-increased ATAC-seq peaks. PWM logos of selected retinal progenitor/neurogenic bHLH TFs are given for comparison. JASPAR IDs of the selected TFs can be found in Methods. **F**. Barchart showing Biological Process (BP) Gene Ontology (GO) term enrichment of genes adjacent to *Crx*^*K88N*^-reduced ATAC-seq peaks.

### *Crx*^*K88N*^-increased accessibility CREs show progenitor cell regulatory signatures and are developmentally silenced during photoreceptor differentiation

Lastly, we asked how CRX K88N ectopic activities interfere with WT CRX functions and lead to dominant photoreceptor deficits that are distinct from all characterized *Crx* animal models (18,22,23). The CRX K88N’s DNA binding preference at Q_50_ HD motifs resembles that of other important retinal HD TFs belonged to the native paired-class Q_50_ HD family (11,40). In the developing mouse retinas, these HD TFs are highly expressed in retinal progenitor cells, coordinate the dynamic chromatin landscape changes during retinal neurogenesis, and are down-regulated in differentiating photoreceptors (41,42). We thus predicted that CRX K88N ectopic activities affect the chromatin landscape changes at subset of progenitor CREs that are usually bound by progenitor Q_50_ homeoproteins.

Using a previously published ATAC-seq dataset of normal mouse retinal development (28), we found that *Crx*^*K88N*^-increased ATAC-seq peaks showed the strongest accessibility at neonatal ages P0 and P3, followed by a gradual decrease in accessibility as photoreceptors undergo differentiation (Figure 4C). Re-analysis of a published VSX2 (Q_50_ HD TF) retinal ChIP-seq data (42) showed that embryonic day 14.5 (E14.5) but not adult VSX2 binding is enriched at the *Crx*^*K88N*^-increased ATAC-seq peaks (Figure 4D). In the embryonic retina, VSX2 is expressed in retinal progenitor cells; in the adult retina, VSX2 expression is maintained in bipolar cells and Müller glia but lost in mature photoreceptors (43-45). Thus, the enrichment of embryonic VSX2 binding and the depletion of adult VSX2 binding at *Crx*^*K88N*^-increased ATAC-seq peaks suggests that CRX K88N ectopic activity is likely impeding the silencing of progenitor programs instead of directing photoreceptor precursors into alternative cell lineage programs. Additionally, *de novo* motif enrichment analysis of sequences under *Crx*^*K88N*^-increased ATAC-seq peaks found patterns characteristic of basic helix-loop-helix (bHLH) neurogenic transcription factor consensuses (Figure 4E), corroborating the potential functionality of *Crx*^*K88N*^-increased ATAC-seq peaks in regulating neurogenic programs during normal development. Indeed, gene ontology (GO) analysis revealed that genes adjacent to *Crx*^*K88N*^-increased ATAC-seq peaks were implicated in general neuronal development (Figure 4F). Collectively, these pieces of evidence suggest that instead of initiating chromatin accessibility *de novo*, CRX K88N is redirected to subset of retinal progenitor CREs enriched for the Q_50_ HD motifs that are regulated by retinal progenitor Q_50_ homeoproteins in normal development. Mutant CRX K88N activities might impede the complete silencing of progenitor regulatory programs and affect development progression in a dominant manner.

### *Crx*^*E80A*^ retinas show opposite chromatin accessibility changes at CREs enriched for monomeric and dimeric K_50_ HD motifs

In contrast to the K88N mutation that drastically alters CRX HD’s DNA binding specificity, E80A does not affect CRX HD’s DNA binding sequence preference *per se* but rather slightly reduces the specificity – binding more promiscuous – at monomeric HD motifs and drives increased expression of target genes in developing photoreceptors (17). Yet, the *Crx*^*E80A/+*^ and *Crx*^*E80A/A*^ retinas are defective in photoreceptor terminal differentiation suggesting additional mechanisms. We thus asked whether CRX E80A’s reduced DNA binding specificity and impaired cooperative dimerization impact chromatin remodeling essential for photoreceptor terminal differentiation. Among the consensus ATAC-seq peaks identified in WT, *Crx*^*E80A/+*^, and *Crx*^*E80A/A*^ retinas, a set of peaks (n=1681) showed significantly increased ATAC-seq signal intensity in the *Crx*^*E80A/A*^ retinas compared to WT, with the *Crx*^*E80A/+*^ retinas showing intermediate intensity at these loci (Figure 5A). The *Crx*^*E80A*^-increased ATAC-seq peaks showed increased CRX E80A binding intensity in the *Crx*^*E80A/A*^ retinas and were enriched for the WT CRX consensus monomeric K_50_ HD motifs (Figure 5B). No additional TF motif was found significantly enriched in these peaks suggesting a major contribution from CRX E80A binding and activity.

**Figure 5.**
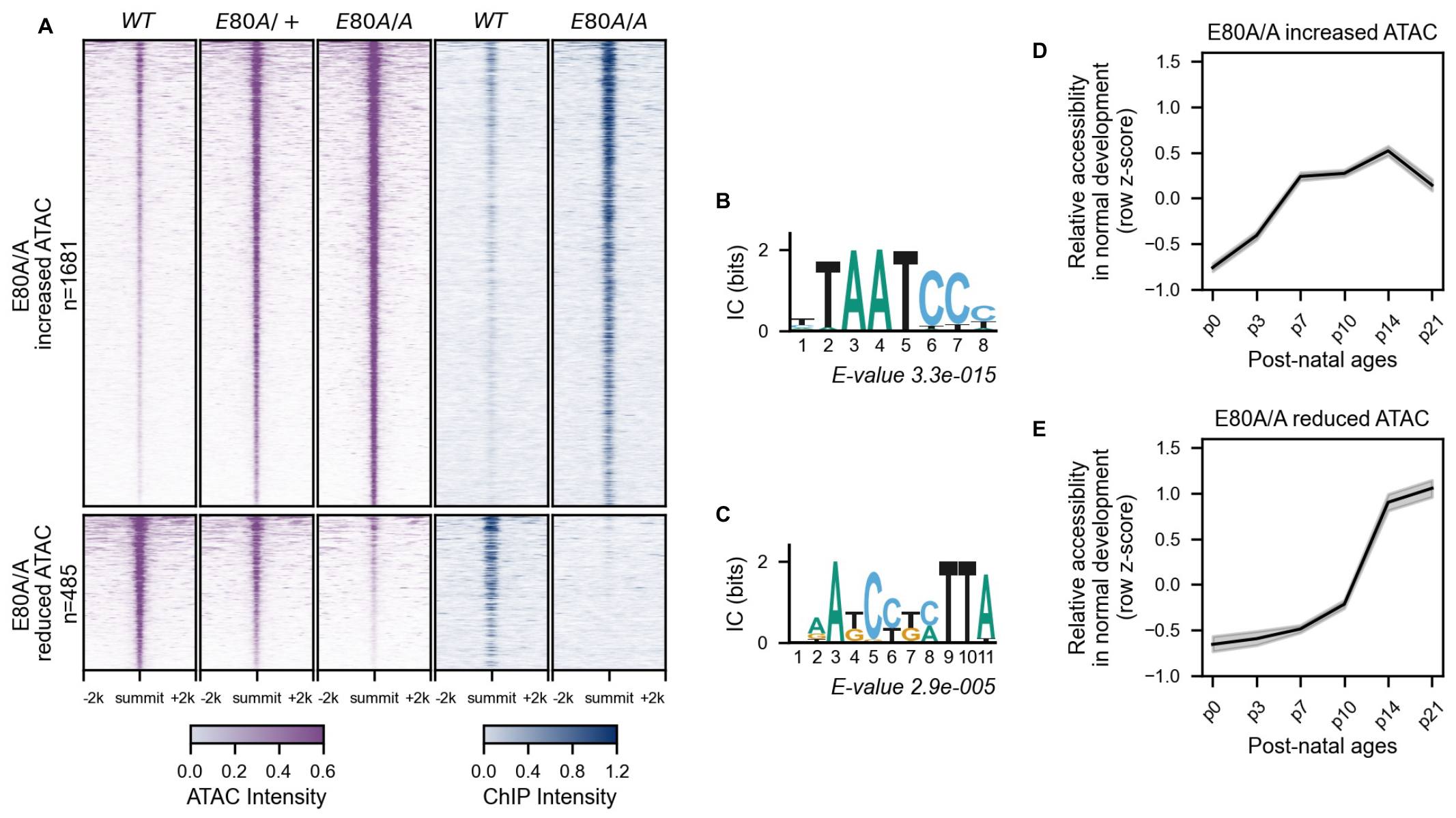
*Crx*^*E80A*^ retinas show defective chromatin remodelling at CREs enriched for dimeric K_50_ HD motifs. **A**. Heatmaps depicting the normalized ATAC-seq or CRX ChIP-seq signal intensities at *Crx*^*E80A*^ differentially accessible ATAC-seq peaks. **B.&C**. PWM logos and enrichment significance *E-values* of STREME *de novo* discovered HD motifs under *Crx*^*E80A*^ differentially accessible ATAC-seq peaks. **D.&E**. Line plots showing the average developmental accessibility kinetics of *Crx*^*E80A*^ differentially accessible ATAC-seq peaks. The developmental ATAC-seq data is from Aldiri et al. 2017.

In contrast, a smaller subset of peaks (n=485) showed significantly decreased ATAC-seq signals in the *Crx*^*E80A/A*^ retinas compared to WT, again with the *Crx*^*E80A/+*^ retinas showing intermediate signal reduction (Figure 5A). The *Crx*^*E80A*^-reduced ATAC-seq peaks showed diminished CRX E80A binding in the *Crx*^*E80A/A*^ retinas likely relate to defective chromatin remodeling at these loci. Interestingly, *de novo* motif searching of sequences under the *Crx*^*E80A*^-reduced ATAC-seq peaks identified a dimeric P3 sequence pattern resembling dimeric K_50_ HD motifs, highlighted by a strong preference of cytosine (C) at the spacer positions 5 and 6 (Figure 5C). This is in line with Coop-seq observation *in vitro* where E80A reduced CRX HD’s cooperative binding at P3 dimeric sequences (Figure 2D). Intriguingly, close examination of the FIMO-identified dimeric K_50_ HD motifs under *Crx*^*E80A*^-reduced ATAC-seq peaks (Figure 5C) found that these dimeric motifs are often consist with two low-affinity half-sites (Figure S3A, Methods). This is distinct from previous analyses that have focused mostly on the highest affinity (not the strongest cooperativity) dimeric K_50_ HD motifs (8,25). This suggests that cooperative dimerization might be critical for WT CRX regulatory activities at specific dimeric K_50_ motifs *in vivo* and in the chromatin context. Thus, impaired cooperative dimerization likely reduces CRX E80A’s stable binding at specific dimeric K_50_ HD motifs which in turn affects the chromatin remodeling activity at CREs where cooperative WT CRX-dimeric K_50_ HD motif interactions are essential.

### *Crx*^*E80A*^-differentially accessible CREs exhibit distinct chromatin remodeling kinetics in normal development

Next, we sought to understand how differential chromatin accessibility relates to the perturbed photoreceptor differentiation in the *Crx*^*E80A/+*^ and *Crx*^*E80A/A*^ retinas. WT CRX activity is required at subset of its binding sites for retinal-specific chromatin remodeling that facilitates post-natal photoreceptor differentiation (36). Using a previously published ATAC-seq dataset of normal mouse retinal development (28), we found that both *Crx*^*E80A*^-increased and *Crx*^*E80A*^-reduced ATAC-seq peaks gain accessibility during post-natal retinal development (Figures 5D-5E). Yet, the two sets of peaks differ in their kinetics of accessibility gain. In WT retinas, the *Crx*^*E80A*^-increased ATAC-seq peaks exhibit a strong increase in accessibility during early photoreceptor development, from post-natal day 0 (P0) to P7, followed by a moderate change from P7 to P21. The *Crx*^*E80A*^-reduced ATAC-seq peaks are characterized by an exponential gain in accessibility between P10 and P14 with smaller changes prior to P10 and after P14. The P10-P14 time points represent a critical window of photoreceptor differentiation characterized by a dramatic change in the photoreceptor transcriptome (46) and the elaboration of Outer Segments (OSs), the subcellular structures where phototransduction in photoreceptors occurs (47). The distinct developmental accessibility kinetics suggest that CRX E80A activity at monomeric and dimeric K_50_ HD motifs might affect gene expression at different stages of photoreceptor development in the mutant mouse retinas.

### Monomeric and dimeric K_50_ HD motifs associated with stage-specific photoreceptor gene mis-expression in the *Crx*^*E80A*^ retinas

To identify the likely direct impacts of CRX E80A mutant activity on gene expression, we focused on the “CRX-dependent activated genes” (CRX-DAGs) defined previously (17). CRX-DAGs are adjacent to at least one CRX binding site and their expressions are significantly reduced in the loss-of-function model *Crx*^*R90W/W*^. CRE-gene association analysis using the ATAC-seq identified CREs that also overlapped with CRX binding revealed that many CRX-DAGs are potentially under combinational regulations of monomeric and dimeric K_50_ HD motifs (Supplementary Table S6, Methods). Specifically, 28.87% of CRX-DAGs are associated with CREs that only contain monomeric K_50_ HD motifs while 61.27% of CRX-DAGs are associated with one or more CREs that contain both monomeric and dimeric K_50_ HD motifs (Figure 6A). This observation raises a possibility that, in addition to high-affinity monomeric motifs (20,26), CRX action on dimeric motifs is required for the regulation of a specific subset of target genes during photoreceptor differentiation. In support of this idea, by analyzing a previously published RNA-seq dataset of normal mouse retinal development (28), we found that genes regulated by both monomeric and dimeric K_50_ HD motifs display a more dramatic increase in expression at photoreceptor terminal differentiation (Figure S4A) coinciding with the exponential gain in accessibility of CREs enriched for dimeric K_50_ HD motifs between P10 to P14 (Figure 5E).

**Figure 6.**
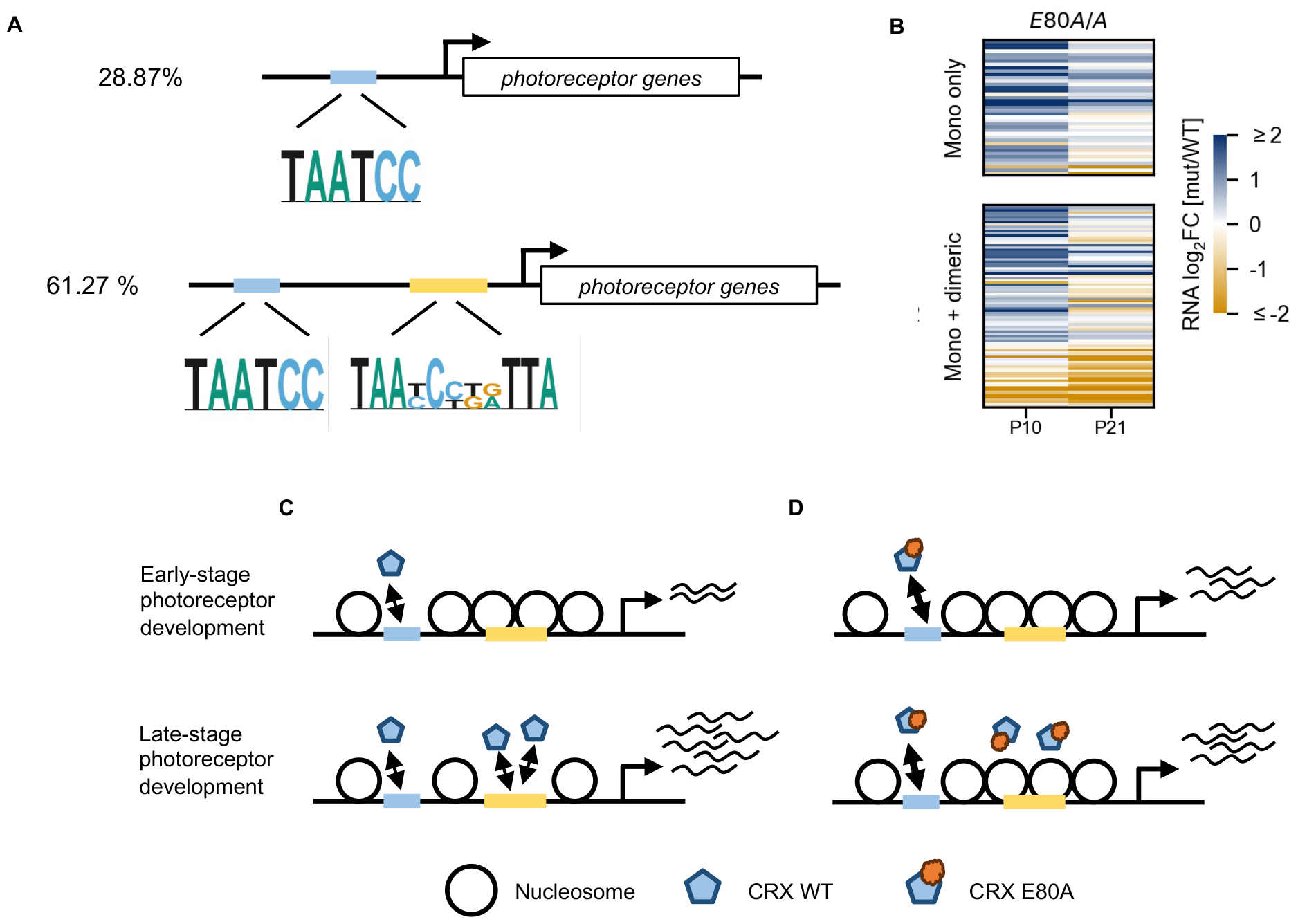
CRX E80A differential activity at monomeric and dimeric K_50_ HD motifs contributes to gene mis-expression at different stages of photoreceptor development. **A**. Diagrams of photoreceptor genes regulated solely by monomeric K_50_ HD motifs (top) or combinationally by monomeric and dimeric K_50_ HD motifs (bottom). For simplicity, representative motif logos are shown. The relative position of the motifs is arbitrary. **B**. Heatmap comparing the CRX-DAG expression changes in the *Crx*^*E80A/A*^ retinas at ages of post-natal day 10 (P10) and day 21 (P21). The gene sets in the heatmaps are as defined in **A. C.&D**. Schematics demonstrating the K_50_ HD division-of-labor model in regulating photoreceptor epigenome and transcriptome at different stages of development.

To understand how altered CRX E80A-DNA interactions at monomeric and dimeric K_50_ HD motifs relate to photoreceptor gene mis-expression in the developing (P10) and mature (P21) *Crx*^*E80A*^ mouse retinas, we reanalyzed the RNA-seq data generated in our previous study (17). In the P10 *Crx*^*E80A/A*^ retinas, CRX-DAGs associated either solely with monomeric K_50_ HD motifs or also with dimeric K_50_ HD motifs showed over-expression compared to WT (Figure 6B, P10 columns). Since regulatory elements enriched for the monomeric K_50_ HD motifs display an early chromatin remodeling profile in normal development (Figure 5D), it is likely that CRX E80A promiscuous binding at monomeric K_50_ HD motifs (17) accelerated the chromatin remodeling and/or directly enhanced the expression of CRX-DAGs in the developing *Crx*^*E80A/A*^ mutant retinas. In the P21 adult *Crx*^*E80A/A*^ retinas, expression of the P10 *Crx*^*E80A/A*^-overexpressed genes showed two patterns: genes associated solely with monomeric K_50_ HD motifs became comparable to WT or remained over-expressed but of a much lower magnitude; genes associated with both monomeric and dimeric K_50_ HD motifs became comparable to WT and even significantly down-regulated (Figure 6B, P21 columns). The *Crx*^*E80A/+*^ retinas showed similar patterns of gene mis-expression but at a less severe degree (Figure S5A). The selective down-regulation of dimeric K_50_ HD motif associated genes may be explained by CRX E80A’s impaired cooperative dimerization and subsequently defective chromatin remodeling at regulatory elements enriched for the dimeric HD motifs. Importantly, genes encoding structural proteins of photoreceptor OSs and molecules involved in the second messenger cascade of the visual cycle are enriched in this specific set of CRX target genes (Figure S5B, Supplementary Table S6). The precise regulation of these genes is fundamental to the integrity of photoreceptor OSs and functions (48). Perturbations of these genes have been associated with inherited retinal dystrophies that affect rods, cones, or both, and cause blindness (49). Thus, under-expression of these genes in the *Crx*^*E80A*^ retinas likely underlie the defective photoreceptor terminal differentiation and functions.

Additionally, a subset of CRX-DAGs associated with dimeric and monomeric K_50_ HD motifs were down-regulated at P10 and became more severely down at P21 (Figures 6B, S5A). Close examination of this gene set revealed that they are implicated in cone photoreceptor structures and functions. This observation suggests the CRX cooperative binding on dimeric K_50_ HD motifs is important for cone gene regulation in development.

### Monomeric and dimeric K_50_ HD motifs demonstrate E80A dosage dependent regulatory activity changes in *ex plant* retinas

The analyses above clearly demonstrate a close relationship between dysregulated CRX E80A activity at different K_50_ HD motifs and gene mis-expression in the *Crx*^*E80A*^ retina. Yet, it remains unclear whether CRX E80A acts directly or through modulating chromatin accessibility to affect gene expression. To evaluate the relative contribution of CRX E80A-DNA interactions to transcription regulation of non-chromatin templates, we selected and tested a representative set of CRX regulated genomic targets in episomal plasmids with Massively Parallel Reporter Assays (MPRAs) in *ex plant* WT and mutant mouse retinas (Figures 7A, S6A-S6B, Methods). Comparison of the MPRA results between genotypes found a CRX E80A dosage-dependent gain in monomeric K_50_ HD motif activity (Figure 7B), suggesting that CRX E80A binding at monomeric motifs can directly drive increased target gene expression in the developing retinas. Different from the monomeric motifs, dimeric K_50_ HD motifs showed similar degree of activity reduction in both *Crx*^*E80A/+*^ and *Crx*^*E80A/A*^ retinas (Figure 7C). This observation suggests that CRX regulatory activity at dimeric K_50_ HD motifs may be sensitive to CRX dosage which is in line with much lower binding affinity of dimeric K_50_ HD motifs *in vitro* compared to the consensus monomeric K_50_ HD motif (8). The MPRA results, combined with alterations of photoreceptor epigenome *in vivo*, suggest that CRX E80A mis-regulates gene expression by acting through both chromatin remodeling and direct transactivation pathways.

**Figure 7.**
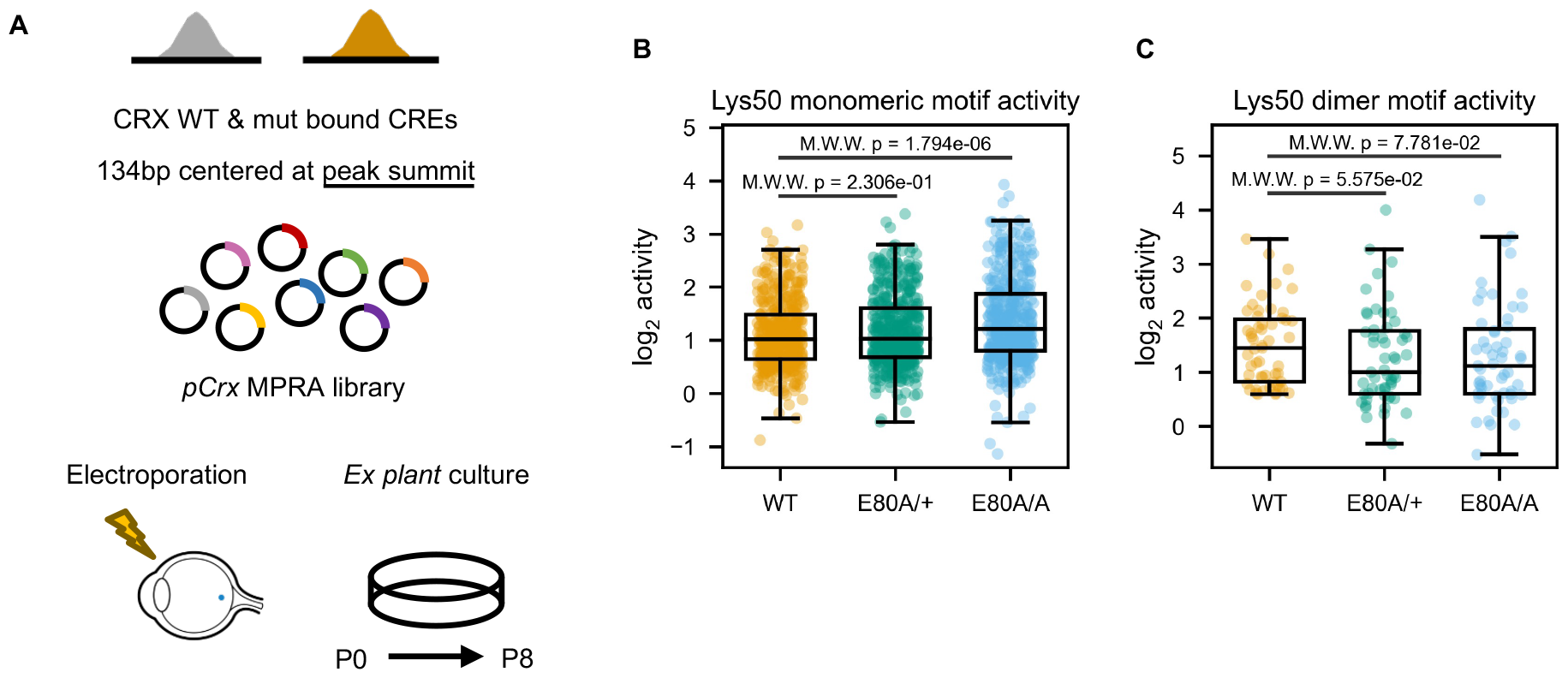
Monomeric and dimeric K_50_ HD motifs associated with E80A dosage dependent activity changes in retinal *ex plant* MPRAs. **A**. Schematics showing *ex plant* retinal MPRA experimental pipeline. **B.&C**. Box and strip plots comparing monomeric (**B**) or dimeric (**C**) K_50_ HD motif activities in *ex plant* cultured WT, *Crx*^*E80A/+*^ and *Crx*^*E80A/A*^ retinas. In panel **B**, CREs overlapped with ATAC-seq peaks that were not significantly reduced in the *Crx*^*E80A*^ retinas are plotted. In panel **C**, CREs overlapped with ATAC-seq peaks that were significantly reduced in the *Crx*^*E80A*^ retinas are plotted. *p-values* of Mann-Whitney-Wilcoxon tests are annotated.

## Discussion

Through molecular characterization of mutant proteins, integrated analysis of the epigenome and CRX regulome in developing mouse retinas, we have advanced our understanding of the distinct pathogenic mechanisms of CRX HD mutations, E80A and K88N, that are associated with dominant CoRD and dominant LCA in humans. Besides the previously identified differential impacts on CRX HD-DNA binding specificity (17), E80A and K88N mutations also differently alter CRX HD’s cooperative dimerization at specific dimeric HD sequences. The mutation specific effects on DNA binding specificity and cooperativity underlie the chromatin landscape changes that explain the distinct dominant photoreceptor gene mis-expression patterns and developmental deficits in the mutation *knock-in* mouse retinas. Despite severely disorganized retinal morphology, no obvious photoreceptor degeneration was observed in either *Crx*^*E80A*^ or *Crx*^*K88N*^ retinas (17), suggesting the molecular changes observed *in vivo* largely attribute to CRX’s intrinsic functional changes. Integrated analysis of the epigenome and transcriptome dynamics in normal development and in the *Crx*^*E80A*^ and *Crx*^*K88N*^ mutant retinas highlight an underappreciated role of DNA-mediated CRX cooperative dimerization in ensuring proper temporal chromatin and gene expression changes at different stages of photoreceptor development.

HD residue 50 (corresponding to CRX K88) determines paired-class HDs’ DNA binding specificity at monomeric and dimeric HD motifs as well as HD’s cooperative binding at palindrome dimeric motifs (8,37,38,50). Consistently, only the K88N mutation, previously found to alter CRX HD’s DNA binding specificity from K_50_ to that resembling a Q_50_ HD (17), drastically enhances CRX HD’s cooperative dimerization at the P3 palindrome-containing *rhodopsin* promoter *BAT-1* probe (Figure 1C). Coop-seq reveals that K88N HD demonstrates strong cooperative binding at all P3 sequences compared to the much weaker and selective cooperativity of WT HD (Figure 2D). The drastic difference between WT and K88N HDs in the scale and spectrum of binding cooperativity agrees with the random-site selection assays (8) where a Q_50_ HD bound strongly to many P3 sequences yielding a 5’-TAATPyNPuATTA-3’ consensus while a K_50_ HD only selected the monomeric consensus 5’-TAATCC-3’ under the same conditions as the Q_50_ HD and three additional rounds of selection were required to recover the dimeric consensus 5’-TAATCCGATTA-3’. Thus, the severe photoreceptor developmental deficits in the *Crx*^*K88N*^ retinas are likely attributed to CRX K88N’s drastically different DNA binding specificity and cooperativity.

In the developing WT mouse retinas, CRX binding at K_50_ HD motifs is essential for photoreceptor differentiation through facilitating chromatin remodeling and regulating target gene expression (36). Since CRX K88N mutant protein prefers a different set of DNA motifs than WT CRX, the diminished activity at canonical CRX motifs leads to defective chromatin remodeling at photoreceptor CREs and loss of target gene expression in the *Crx*^*K88N/N*^ retinas (17). Yet, the dominant inheritance pattern and the more severe gene mis-expression in the *Crx*^*K88N/N*^ retinas than the loss-of-function *Crx*^*R90W/W*^ retinas (Figure 3E) suggests that CRX K88N ectopic activities affect other regulatory programs in addition to those regulated by WT CRX. Indeed, subset of regulatory elements enriched for the Q_50_ HD motifs failed to be silenced in the *Crx*^*K88N/+*^ and *Crx*^*K88N/N*^ mutant retinas (Figure 4A). The silencing defect is not observed in the *Crx*^*R90W/W*^ retinas suggesting loss of WT functions alone is an insufficient explanation. Although one may predict novel heterodimeric CRX WT-K88N cooperative binding and ectopic activities, the same set of ATAC-seq peaks were found differentially accessible in the *Crx*^*K88N/+*^ retinas (Figure 4A). A separate *de novo* motif analysis using *Crx*^*K88N/+*^ differentially increased ATAC-seq peaks revealed a similar dimeric Q_50_ HD-like motif to that identified using *Crx*^*K88N/N*^ data (Figure 4B, Supplementary Table S4) instead of a composite K_50_-Q_50_ HD motif. In addition, structural and biochemical studies have demonstrated that paired-class HD cooperative dimerization is mediated through specific HD-DNA interactions (8,51). Thus, the divergent DNA binding specificity of K_50_ and Q_50_ HDs likely restrains the formation of CRX WT-K88N heterodimeric complexes on the majority of dimeric K_50_ or Q_50_ HD motifs.

In normal development, many HD containing TFs expressed in the retinal progenitor cells belong to the Q_50_ HD paired-class subfamily, including many eye field TFs (11,40,52). The expression and functions of progenitor Q_50_ HD TFs promotes retinal progenitor cell proliferation and thus is inhibitory to neurogenesis and subsequently post-mitotic precursor cell differentiation (44,53,54). In post-mitotic photoreceptor precursors, these progenitor Q_50_ HD TFs are down-regulated accompanying the silencing of proliferative programs while the expression of K_50_ HD TFs, including OTX2 and CRX, activates photoreceptor differentiation programs (55-58). It is conceivable that there is a brief period in early photoreceptor precursor cells when the progenitor Q_50_ HD TF regulated chromatin regions are not completely silenced while CRX proteins are already present (Figure 8). In the *Crx*^*K88N/+*^ and *Crx*^*K88N/N*^ mutant retinas, CRX K88N proteins likely bind to subset of these progenitor CREs, impede the complete silencing of proliferative programs, and consequently interfere with the entry into photoreceptor differentiation programs. In addition, a *CRX* p.Lys88Glu (K88Q) mutation has been reported in an individual with dominant LCA (59) supporting our model that the resemblance of N_50_ and Q_50_ HD DNA binding specificity and cooperativity underlies the severe, dominant photoreceptor developmental deficits in the *Crx*^*K88N*^ mouse models. Alternatively, it is also possible that CRX K88N ectopic activities simply create “regulatory noises” that affect the fidelity of photoreceptor developmental programs. In summary, CRX target specificity is not only critical for activating photoreceptor developmental programs but also crucial for the proper silencing of early programs that could be inhibitory to photoreceptor differentiation at late stages.

**Figure 8.**
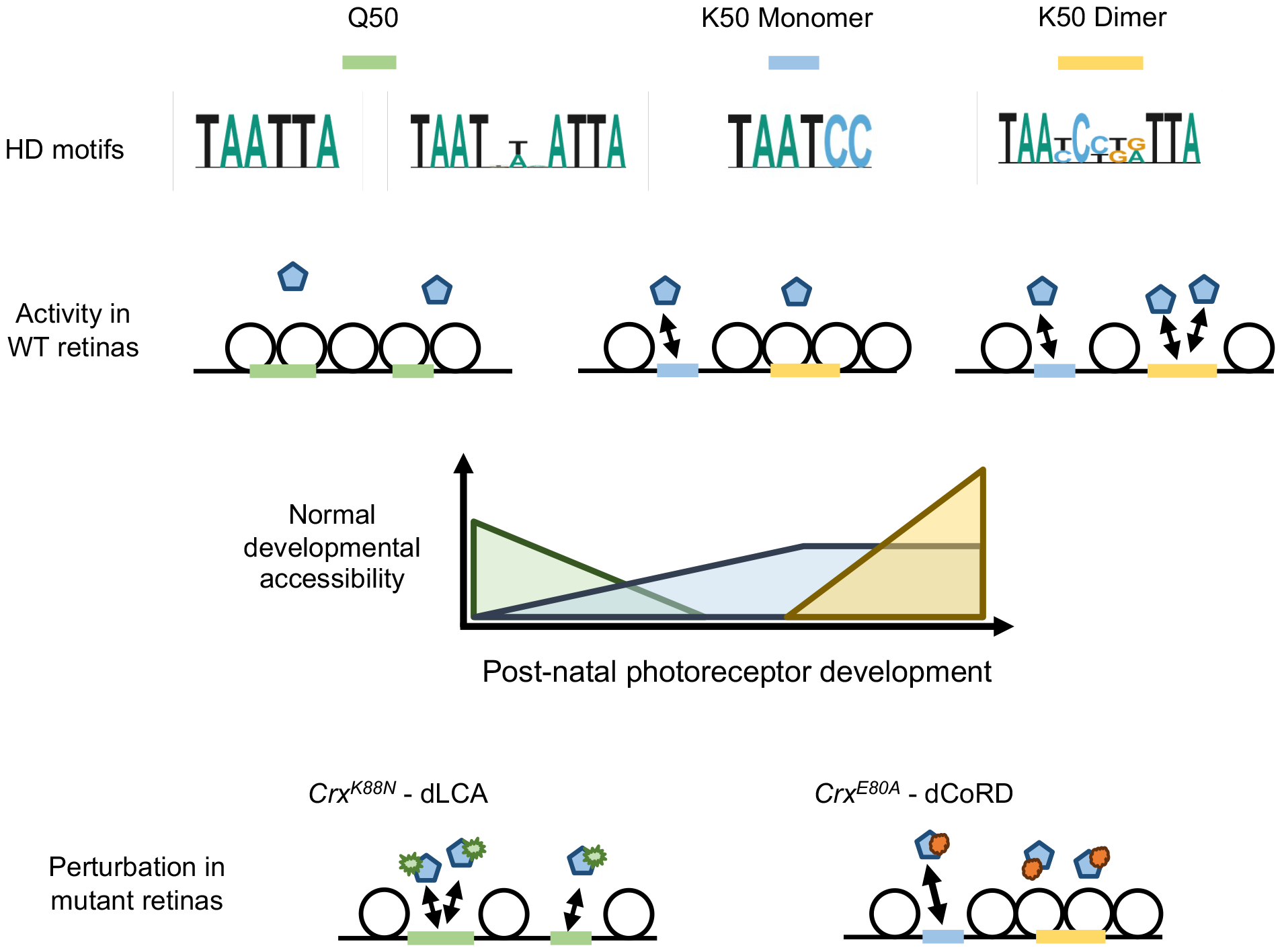
Model schematics.

Different from K88N’s global impacts on CRX HD-DNA interactions, E80A mutation increases HD’s promiscuous binding at monomeric K_50_ HD motifs (17) but reduces cooperative binding at dimeric K_50_ HD motifs (Figure 2D). CRX E80A’s differential interactions with K_50_ HD motif sub-types parallel the hyper-activation of subset photoreceptor genes in developing *Crx*^*E80A*^ retinas and the hypo-activation of the same set of genes in adult *Crx*^*E80A*^ retinas (17). These observations point to a K_50_ HD motif division-of-labor model where early-stage photoreceptor development is mainly mediated by CRX interactions with monomeric K_50_ HD motifs, while photoreceptor terminal differentiation additionally relies on interactions with dimeric K_50_ HD motifs that specifically requires cooperative dimerization (Figure 8). This dichotomy of K_50_ HD motif usage likely relates to the different CRX-chromatin interaction kinetics that is determined by the underlying DNA sequences and TF concentrations in developing photoreceptors (Figures 5D-5E, S4B-S4C). Specifically, CRX binds monomeric K_50_ HD consensus motifs with high affinity which can drive strong gene activity at relatively low CRX concentration, such as in photoreceptor precursors (Figures S4B-S4C). However, CRX binding also easily saturates at high affinity monomeric motifs resulting in a small dynamic range of regulatory activity. In contrast, CRX dimeric binding to K_50_ HD P3 sequences is of much lower affinity and requires cooperative dimerization (8). These properties render dimeric K_50_ HD motifs less active at low CRX concentration and remain responsive to higher and a wider range of CRX concentrations (Figures S4B-S4C). Many CRX-DAGs that are regulated by monomeric and dimeric K_50_ HD motifs encode proteins in the phototransduction pathway. Proteins in this pathway need to express robustly and remain dynamic in response to changes in ambient illumination. Thus, it is likely that combinational regulation of monomeric and dimeric K_50_ HD motifs imparts functional specification of CRX to regulate a subset of target genes that play specific physiological roles in photoreceptor biology. Collectively, precision in CRX-DNA interactions is important for not only the quantitative regulation but also the temporal regulation of photoreceptor gene expression.

Although the K_50_ HD motif division-of-labor model explains the general CRX target gene expression alternations in the *Crx*^*E80A*^ retinas, it remains unclear why the subset of CRX dependent genes implicated in cone photoreceptor development and functions is down-regulated in both differentiating and mature mutant retinas (17) (Figures 6B, S5A) . While cone photoreceptors were born in these mutant retinas, the loss of cell-type specific gene expression in early post-natal development suggests defective differentiation (17). One possibility is that cone genes are more dependent on CRX activity at dimeric K_50_ HD motifs. In support of this prediction, most of the cone-enriched CRX-dependent activated genes (81.25%) are associated with dimeric K_50_ HD motifs as compared to only 62.32% of rod-enriched genes in the same category (Figure S4D). An alternative but not mutually exclusive model is that cones are more sensitive to perturbations in CRX activity. In the developing and mature mouse retinas, cones are dependent on a different repertoire of TFs than rods for differentiation and functions (47,60-63). Many rod-specific transcription factors collaborate with CRX in strongly activating rod gene expressions. Small perturbations in CRX activity might thus be dynamically compensated by CRX collaborating factors. In contrast, although many nuclear receptor family TFs have been identified to mediate M-vs S-cone subtype differentiation, these factors are dispensable for cone cell genesis, development, or survival in early post-natal ages (60). It is possible that CRX plays a major role in regulating general cone cell development and functions. The mouse retina is rod-dominant and thus is limited in resolution of cone-related mechanistic understandings. Quantitative characterization of CRX molecular functions in a pure cone population and comparison with the *Crx*^*E80A*^ model warrants further study to elucidate regulatory principles in early photoreceptor development and in *CRX*-linked dominant CoRD.

Collectively, our findings support a unifying model in which differential CRX interactions with different sub-types of HD motifs underlie cell-stage specific chromatin remodeling and temporal gene regulation during photoreceptor development. Disease-associated mutations in CRX can be classified into two major groups – frameshift/nonsense mutations in the transcription effector domain and missense mutations in the homeodomain. Prior biochemical and mouse model studies of frameshift mutations and the loss-of-function R90W mutation have established that CRX binding and transactivation activity at high affinity monomeric K_50_ HD motifs is essential for photoreceptor gene regulation (18,23,26,64,65). Yet, these mutations either abolish CRX DNA binding or abrogate CRX transactivation activity and lead to rapid photoreceptor degeneration, obscuring the resolution of differential CRX interactions with HD motif sub-types. In this study, we demonstrate that missense mutations in the CRX HD, by two combinational gain- and loss-of-function mechanisms, alter CRX DNA-binding specificity and cooperativity at distinct types of HD motifs. These biochemical property changes result in mutation specific chromatin remodeling defects and impaired photoreceptor differentiation programs in the mutant mouse retinas.

Comparative analysis of the WT and mutant mouse retina epigenome and transcriptome suggests two regulatory principles that coincide with important developmental transitions in photoreceptors (Figure 8). First, the transition from proliferating retinal progenitor cells to committed photoreceptor cells is accompanied by a shift from highly expressed Q_50_ to K_50_ paired-class HD TFs and from regulatory elements enriched with Q_50_ to those with high affinity monomeric K_50_ HD motifs. Second, the transition from early to late-stage photoreceptor development requires the utilization of dimeric K_50_ HD motifs in addition to monomeric K_50_ HD motifs. The first transition involves a sharp change in DNA binding specificity which likely confers sensitivity in newly post-mitotic photoreceptor precursors to respond to low concentration of CRX and quickly fix to a committed photoreceptor precursor status. This strategy may also ensure the rapid elimination of progenitor epigenetic features due to a lack of interacting TFs. In contrast, the second principle imply a functional specialization of CRX at a specific set of target genes whose expression need to remain dynamic and robust in the mature photoreceptors.

Our study here refines the CRX mechanistic model in photoreceptor development, expands our knowledge on the diverse mechanisms that CRX mutations lead to severe, dominant retinopathies, and lays the foundation for future development of therapeutic strategies targeting different pathogenic mechanisms. Our CRX mechanistic model emphasizes the importance of considering interactions between coding CRX variants and non-coding variants in CRX binding sites in modifying clinical phenotypes. Our study also demonstrates, in addition to its unique biochemical properties (8), that paired-class HD cooperative dimerization plays a crucial role in development and its dysregulation can lead to distinct human diseases. The adaption of Coop-seq enables the unbiased identification of CRX HD cooperative dimerization as opposed to non-cooperative co-binding, which are not easily separable in selection-based TF-DNA binding assays. Our study provides additional hints towards understanding the structural basis of paired-class HD cooperative dimerization on palindrome DNA sequences (51). Multiple missense mutations at the CRX E80 residue have been reported in dominant CoRD cases, including p.E80K (Clin Var VCV000099599), p.E80G (VCV000865803), and p.E80A (VCV000007416). Systematic investigation on how disease-associated variants affect CRX HD-DNA contacts and/or intramolecular contacts with other HD residues would guarantee new structural insights. Lastly, given that HD TF molecular mechanisms of action are conserved across evolution and in different tissues and organs, we envision our CRX study will also shed light on the study of other homeoproteins that hopeful lead to advance in medicine for the associated diseases.

## Resource availability

### Lead contact

Further information and requests for resources and reagents should be directed to and will be fulfilled by the lead contact Shiming Chen.

#### Materials Availability

- All unique/stable reagents generated in this study are available from the lead contact with a completed materials transfer agreement.

#### Data and Code Availability

- The data underlying this article are available in the NCBI Gene Expression Omnibus (GEO) database at https://www.ncbi.nlm.nih.gov/geo/.
- Customized scripts and any additional information required to reproduce the analysis in this paper will be available from GitHub at https://github.com/YiqiaoZHENG/CRXHD_epigenome.git (data visualization) and https://github.com/YiqiaoZHENG/CRXHD_mpra.git (dedicated MPRA design, sequencing library preparation, and data analysis).

### Methods

#### Animal study and sample collection

#### Ethics statement

All procedures involving mice were approved by the Animal Studies Committee of Washington University in St. Louis and performed under Protocol 21-0414 (to SC). Experiments were carried out in strict accordance with recommendations in the Guide for the Care and Use of Laboratory Animals of the National Institutes of Health (Bethesda, MD), the Washington University Policy on the Use of Animals in Research, and the Guidelines for the Use of Animals in Visual Research of the Association for Research in Ophthalmology and Visual Sciences. Every effort was made to minimize the animals’ suffering, anxiety, and discomfort.

#### ATAC-seq sample collection and library preparation

The assay for transposase-accessible chromatin with sequencing was performed as previously published (24). Briefly, for each genotype, three biological replicates, two retinas per replicate from one male and one female were pooled. The pooled retinas were washed twice with cold PBS before being dissociation in 250μl TESCA buffer (50 mM TES, 0.36 mM calcium chloride, pH 7.4 at 37°C) containing 2% collagenase (Sigma) for 10mins at 37°C. The dissociation mixture was subjected to DNase I treatment (10ul DNaseI at 2U/μl, New England Biolabs) for an additional 3mins at 37°C. The DNase I activity was quenched by adding 500ul DMEM (Gibco™) with 10% HI-FBS (Gibco™) and incubating for 5mins at room temperature. The reaction mixture was passed through a 40μm strainer (Falcon® Corning) twice and centrifugated at 400rcf for 5mins at 4°C to collect cells. The cells were washed twice with cold PBS, centrifugated at 500rcf for 5mins at 4°C, and gently resuspended in 400μl cold lysis buffer (ATAC-RSB, 0.1%Tween20, 0.1%NP-40). The lysis reaction was carried out on ice for 5mins. The nuclei were separated from the lysis mixture by centrifugation at 500rcf for 5mins at 4°C, washed once with cold nucleus resuspension buffer (ATAC-RSB, 0.1%Tween20, 0.01% Digitonin), and collected by centrifugation at 500rcf for 10mins at 4°C. The nuclei were gently resuspended in 20μl 2x TD (Illumina Cat #20034211) buffer and counted with a hemocytometer by staining the intact nuclei with SYBR® Green I Nucleic Acid Stain (Invitrogen™). 10k nuclei were aliquoted and subjected to tagmentation (50 μl reaction with 2.5μl Illumina Tagment DNA TDE1 Enzyme) for 30mins at 37°C following a standard protocol. The recipe for buffer ATAC-RSB is 10mM Tris-HCl, 10mM NaCl, 3mM MgCl_2_.

The tagmented DNAs were cleaned up by the MinElute® PCR purification kit (QIAGEN), eluted in 23μl Nuclease-Free Water (Invitrogen™ Ambion®) and stored at -20°C if not immediately used for library preparation. The ATAC-seq libraries were constructed by PCR amplification of the tagmented DNAs for 10 cycles using the NEBNext® High-Fidelity 2X PCR Master Mix (New England Biolabs). The PCR reactions were directly subjected to double size selection (0.55x and 1.55x) with the AMPure XP Beads (Beckman Coulter) following the manufacturer’s protocol. The cleaned-up ATAC-seq fragments were eluted with 12μl Nuclease-Free Water (Invitrogen™). The quantity and quality of the ATAC-seq libraries was assayed using the Qubit™ 3 Fluorometer (Invitrogen™) and the Bioanalyzer (Agilent, Santa Clara, CA) prior to sequencing.

#### MPRA plasmid library construction

The MPRA plasmid library was constructed following published protocols (25). Briefly, the library of 200mer oligonucleotides was ordered directly from Twist Bioscience (South San Francisco, CA). Each oligo contains a 134bp testing cis-regulatory element (CRE) sequence and a unique 10bp barcode flanked by sequences required for cloning (Figure S6A). The plasmid library was amplified with NEB Phusion® High-Fidelity PCR Master Mix with HF Buffer (New England Biolabs). The amplified fragments were cloned into the EcoRI-EagI sites of the vector pJK03 (Addgene ID173490). Lastly, the *Crx* promoter-DsRed fragment was amplified from the pCrx-DsRed plasmid (25) and cloned into the SpeI-SphI sites located between the CRE sequence and the barcode. The constructed MPRA plasmid library was amplified by transformation into the NEB® 5-alpha Competent E. coli (High Efficiency) DH5α (New England Biolabs) according to manufacturer instructions. Prior to retinal electroporation, the fragments containing the 10bp unique barcode were amplified from the plasmid library and subjected to Sanger sequencing and 2x150bp Nova-seq to ensure successful library construction. The complete CRE sequences and annotations can be found in the GitHub repository for this paper.

#### MPRA plasmid preparation for electroporation

For each electroporation, 30μg of MPRA plasmid library DNA per retina (3 retinas per replicate, a total of 3-4 replicates per genotype) was aliquoted and brought to 150μl with water. 15μl NaOAc (pH5.2) was added to the DNA followed by vortex. 450μl absolute ethanol was added to the mixture followed by vortex. The DNA was precipitated by centrifugation at 13500rpm for 30mins at 4°C followed by a second wash with 400μl 70% ethanol and centrifugation 13500rpm for 15mins at 4°C. The DNA pellet was then air-dry until semi-transparent and resuspended thoroughly in appropriate amount of PBS. The concentration of the prepared DNA was measured with the Qubit™ 3 Fluorometer (Invitrogen™) and adjusted with PBS to a final concentration of 0.5μg/μl.

#### Retinal ex plant electroporation

Retinas were dissected in DMEM/F12 1:1 buffer (Gibco™) from newborn (P0) WT and *Crx* mutant mice. For each replicate, three retinas were transferred into the electroporation chamber (CUY520P5, NEPA GENE Co. Ltd) filled with the prepared MPRA plasmid DNA solution. The retinas were electroporated with the Electro Square Porator™ (ECM®830, BTX) with settings: LV, V: 30 Volts, Pulse Length: 50 msec, # Pulses: 5, Interval: 950 msec. The electroporated retinas were washed once in the retinal *ex plant* culture medium, carefully transferred onto a 0.2μm 25mm Nuclepore™ Track-Etch Membrane (Whatman®) and cultured for 8 days in the incubator (37°C, 5% CO_2_) with the retinal *ex plant* culture medium (DMEM/F12 1:1, 10% HI-FBS, 1xPenicillin-Streptomycin). The day-8 *ex plant* retinas were collected, washed once in cold PBS, and stored in 30μl Nuclease-Free Water (Invitrogen™) at -80°C before ready for TRIzol extraction.

#### Retinal ex plant RNA/DNA Trizol extraction and purification

The RNA and DNA was extracted from *ex plant* cultured retinas using TRIzol™ Reagent (Invitrogen™) following a modified protocol based on the manufacturer’s protocol. The retinas were thawed on ice, washed once with ice-cold Molecular Biology Grade water (Corning), and resuspended in 400μl TRIzol™ Reagent. The retinas were homogenized with a PRO250 Homogenizer (PRO Scientific Inc.) and a 5mm X 75mm Flat Bottom Generator Probe (PRO Scientific Inc.) at intensity level 3 for 15sec. The homogenized tissue was incubated for 5mins at room temperature before 80μl chloroform was added followed by vigorous mixing by hand for 15sec. The TRIzol-chloroform mixture was incubated at room temperature for 2-3 mins and centrifuged at 12000g at 4°C for 15mins. The top aqueous phase containing RNA was transferred to a new 1.5ml Eppendorf tube. The RNA was purified and concentrated with TURBO DNA-free™ Kit (Invitrogen™) and Zymo RNA Clean & Concentrator™-5 kit (Zymo Research) following manufacturers’ protocols and eluted twice with 15μl Nuclease-Free Water (Invitrogen™).

To extract DNA, any remaining aqueous layer containing RNA was removed from the TRIzol mixture. 200μl DNA back extraction buffer (4 M guanidine thiocyanate, 50 mM sodium citrate and 1 M Tris (free base), 0.5 vol of starting TRIzol Reagent used) was added to the remaining TRIzol mixture, mixed by inversion, and incubated at room temperature for 10mins. The mixture was centrifuged at 12000g at room temperature for 30mins. The top aqueous phase containing DNA was transferred to a new 1.5 ml Eppendorf tube. 160μl (0.4 vol of starting TRIzol Reagent used) of absolute isopropanol was added to the aqueous layer, mixed by inversion, and incubated at room temperature for 5mins before centrifugation at 12000g at 4°C for 15mins to pellet the DNA. The DNA pellet was washed twice with 200μl 70% ethanol and centrifuged at 12000g at 4°C for 15mins. The DNA pellet was then air-dry until semi-transparent and resuspended thoroughly in 30μl of Nuclease-Free Water (Invitrogen™). The samples were left at 4°C overnight for the DNA to fully dissolve. The quantity and quality of the prepared RNA and DNA was determined with the Qubit™ 3 Fluorometer (Invitrogen™).

#### MPRA sequencing library preparation

For each sample, 1μg of purified total RNA was used for cDNA synthesis in 20μl volume using the iScript™ cDNA Synthesis Kit (Bio-Rad Laboratories) following the manufacturer’s protocol. Estimation of the average MPRA library activity was determined by qRT-PCR quantification against the DsRed sequence in MPRA library oligos and the endogenous *Crx* and *Rho* genes (Supplementary Table S1). Three rounds of PCR amplifications were performed to add internal phasing indexes and unique-dual-indexes (UDIs) for Illumina sequencing. PCR1: MPRA library fragments containing the 10bp barcodes were amplified from the cDNA mixture with the Q5® High-Fidelity 2X Master Mix (New England Biolabs). For each sample, 4-8 PCR1 replicates were pooled and purified with Monarch® PCR & DNA Cleanup Kit (New England Biolabs), eluted in 20μl of Nuclease-Free Water (Invitrogen™), and quantified with the Qubit™ 3 Fluorometer (Invitrogen™). PCR2: P1 adapters containing the inner phasing index were added to the amplified fragments from PCR1. PCR3: For each sample, 2μl of PCR2 reaction was used for PCR3 to add the Illumina sequencing adapters and indexes. The final reactions were cleaned up with Monarch® PCR & DNA Cleanup Kit (New England Biolabs), eluted in 30μl of Nuclease-Free Water (Invitrogen™), and quantified with the Qubit™ 3 Fluorometer (Invitrogen™). The sequences of the internal phasing indexes, PCR primers and PCR protocols for MPRA sequencing library preparation can be found in the GitHub repository of this paper.

### Biochemistry

#### Protein expression for EMSA

GST-WT, E80A, K88N and R90W HD peptides were expressed in *E. coli* BL-21 (DE3) cells and purified with GST Spintrap™ columns (Cytiva, Marlborough, MA) as published previously (17,26). The peptides were eluted following the manufacturer’s protocol and buffer exchanged into 1x CRX binding buffer (60mM KCl, 25mM HEPES, 5% glycerol, 1mM DTT) using Amicon centrifugal filters (MilliporeSigma, Burlington, MA). The protein stock was supplemented with 10% glycerol before aliquoted and stored at -80°C.

#### EMSA BAT-1 probe and Coop-seq library synthesis and purification

The *BAT-1* templates (Supplementary Table S1) and IRDye700-labeled reverse complements were ordered directly from Integrated DNA Technologies (IDT). Equal amounts of two oligo strands were mixed in 20μl EB buffer (QIAGEN) and heated to 94°C for 2mins followed by gradual cooling. The Coop-seq libraries were prepared following the Spec-seq library preparation protocol as previously described (17). Briefly, single-stranded Coop-seq library templates (Supplementary Table S1) and IRDye800-labeled reverse complement primers were ordered directly from Integrated DNA Technologies (IDT). 100pmol of template oligos and 125pmol of reverse complement primers F1 were mixed in Phusion® High-Fidelity PCR Master Mix (New England Biolabs). A 15s denaturing at 95°C following a 10-minute extension at 52°C afforded duplex DNAs. The mixture was treated with 1μl Exonuclease I (New England Biolabs) at 37°C for 30mins to remove residual ssDNA. The probes were purified with the MinElute® PCR purification kit (QIAGEN), eluted in an appropriate amount of Nuclease-Free Water (Invitrogen™), and stored at -20°C.

#### EMSA and sample preparation for sequencing

The HD-DNA binding reactions and EMSAs were performed as previously described (17). Briefly, the protein-DNA binding reactions was performed in 1x CRX binding buffer (60mM KCl, 25mM HEPES, 5% glycerol, 1mM DTT). A fixed amount (Supplementary Table S1) of IRDye-labelled BAT-1 probes or Coop-seq libraries were incubated on ice for 30 minutes with varying concentrations of CRX HD peptides in 20μl reaction volume. The reaction mixtures were then run at 4°C in native 12% Tris-Glycine PAGE gel (Invitrogen™) at 160V for 40min. The IRDye-labeled DNA fragments were visualized by Odyssey® CLx and Fc Imaging Systems (LI-COR, Inc.). In the HD-*BAT-1* binding reactions, for visualization purpose, GST tags were cleaved by treating the GST-HD recombinant peptides with 1μl Thrombin protease (1:10 dilution in PBS, Millipore Sigma) for 15mins at room temperature. The GST-HD digestion reactions were used directly in HD-DNA binding assays.

For Coop-seq libraries, the visible bands were excised from the gels. The DNAs were extracted with acrylamide extraction buffer (100mM NH_4_OAc, 10mM Mg(OAc)_2_, 0.1% SDS) and purified with MinElute® PCR Purification Kit (QIAGEN). The extracted Coop-seq DNAs were amplified, barcoded by indexed Illumina primers, then pooled and sequenced on a single 1x50bp Miseq run at DNA Sequencing Innovation Lab at the Center for Genome Sciences & Systems Biology (CGS&SB, WashU).

#### qRT-PCR

Whole retina RNA extraction, purification and cDNA synthesis was performed as previously described (17). Primers used in this study are listed in Supplementary Table S1. qRT-PCR reactions were assembled using the SsoFast™ EvaGreen® Supermix with Low ROX (Bio-Rad Laboratories) following manufacturer’s protocol. Data was obtained from Bio-Rad CFX96 Thermal Cycler (Bio-Rad Laboratories) following a three-step protocol: 1 cycle of 95°C 3 min, 40 cycles of 95°C 10 sec and 60°C 30 sec. Data was exported and further processed with customized python script.

#### BAT-1 variant luciferase reporter assay vector construction

For each *BAT-1* variant, three pairs of oligos (Supplementary Table S1) were designed with each pair containing a four base pair overlap with either the backbone vector or to an adjacent annealed oligo pair. For instance, F1-R1 pairs overlap with the vector and F2-R2 on either end; F2-R2 pairs overlap with F1-R1 and F3-R3 on either end; F3-R3 pairs overlap with F2-R2 and the vector on either end. The oligos were ordered directly from Integrated DNA Technologies (IDT). Each pair of oligos were annealed and the annealed oligo pairs were then ligated and cloned into the pGL2-3xBAT1-luc backbone vector between the restriction sites NheI and XhoI.

#### Cell line transient transfection luciferase reporter assays

HEK293T cell luciferase reporter assays were performed as previously described (17). Briefly, cells were transfected following the calcium phosphate transfection protocol in 6-well plates. Experimental plasmids and usage amounts are described in Supplementary Table 1. 48 hours after transfection, cells were harvested, digested, and assayed for luciferase activity using the Dual-Luciferase Reporter Assay System (Promega) following the manufacturer’s protocol. Data were collected with a TD-20/20 Luminometer (Turner Designs) and processed with customized python scripts.

The HEK293T cells (CRL-3216™) were obtained directly from ATCC (American Type Culture Collection) and used within one year of purchase. The cells were tested negative for mycoplasma contamination.

### Data analysis

#### Determination of relative cooperativity with Coop-seq

For a combinatorial interaction of two (or more) protein species *X* and *Y*, and a particular DNA sequence, *D*_*xx′*_, which is a combination of two half-sites, *x* and *x′*with a spacer, *z*, between them, the dimeric binding interaction can be diagrammed as:

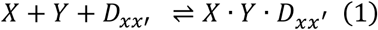

where *X* · *Y* · *D*_*xx′*_ refers to the dimeric protein-DNA complex. Specifically, in our study, we are interested in single protein species *P* binding to a dimeric DNA sequence *D*_*xx’*_. We can rewrite the equation above as:

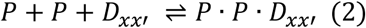

In addition, in our system, the HD peptides (protein species *P*) is also competent to bind each half-site as a monomeric protein-DNA complex. There are two additional states of the DNA sequence being bound:

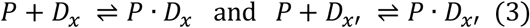

At equilibrium, the probability of a DNA molecule being in the unbound (U), monomerically bound (B_m_), or dimerically bound (B_d_) states (Figure 2B) are:

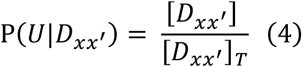

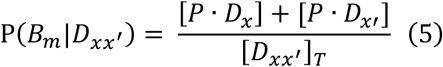

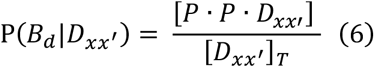

where […] refers to concentrations and [*D*_*xx′*_]_*T*_ as the sum of all the states. The affinity of the protein *P* to a sequence *D*_*x*_ is defined as the association constant *K*_*A*_. The ratios of the probabilities above are related to the association constants as:

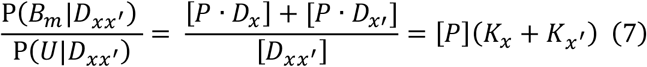

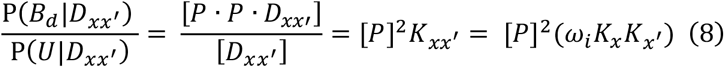

where *ω*_*i*_ is the cooperativity and is unique to each dimeric sequence *D*_*xx′*_ with different half-site combination and spacer. The cooperativity for each unique *D*_*xx′*_ can be written as:

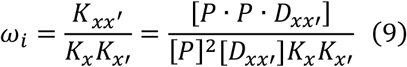

To obtain relative cooperativities relative to some reference sequence *D*_*rr′*_:

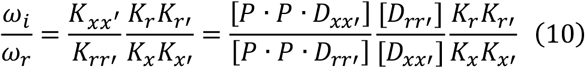

Since we are interested in comparing the cooperativity index *ω*_*i*_ for half-site matched dimeric sequences with a 3bp (P3) *versus* 5bp (P5) spacer (Figures 2C-2D), the 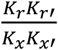 term can be neglected assuming the monomeric binding to half-sites are the same for the two sequence configurations. In a binding reaction involving TF and a library of DNAs, the concentration of the bound and the unbound species are directly proportional to the number of individual DNA molecules in each fraction obtained directly from the sequencing data. With sequencing counts in each fraction, we can accurately estimate the ratios of concentrations from counts with the relationship:

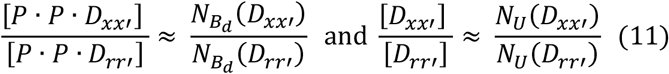

where *N*_*U*_ denotes counts in the unbound fraction and 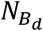 denotes counts in the dimerically bound fraction. We can then rewrite equation 10 as:

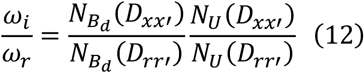

#### Coop-seq data analysis and cooperativity index calculation

The raw sequencing data for monomer and unbound (free) bands were included in our previous publication under GEO accession number GSE223658. The sequencing results were filtered and converted to count ratios following the Spec-seq analysis pipeline previously described (17). Briefly, reads with any mismatch in the conserved regions were discarded and sequences with less than 50 raw read counts were discarded. The relative cooperativity index *ω*_*i*_ for each dimeric sequence was calculated following Equation 12. In Figure 2D, pair-wise relative cooperativity of half-site matched P3-P5 library members is presented. The relative cooperativity was ordered by hierarchical clustering using the python package scipy (v1.11.2) with linkage method “complete” and distance metric “euclidean”. In Figures S2A-S2D, a single reference sequence 5’-TAATGCGCTATTA-3’ was used to calculate relative cooperativity. We chose this sequence based on previous Spec-seq results that WT, E80A and K88N HDs all bind with reasonable affinity to either half-site. The full relative cooperativity index table for all library members can be found in Supplementary Table S2. The P3-P5 matched relative cooperativity index table sorted as in Figure 2D can be found in Supplementary Table S3.

#### ATAC-seq data analysis

2x150bp reads from Illumina NovaSeq2000 were obtained for all samples with an average depth of 54M reads at Genome Technology Access Center at the McDonnell Genome Institute (GTAC@MGI, WashU). For each sample, reads were first run through Trim Galore (v0.6.1) to remove adapter sequences and then QC by FastQC (v0.11.7). The trimmed reads were then mapped to the mm10 genome using Bowtie2 (v2.3.5) with parameters -X 2000 --very-sensitive. Only uniquely mapped and properly paired reads were retained with samtools (v1.12) with parameters -f 0x2 -q 30. Mitochondria reads were removed with samtools (v1.12). Duplicated reads were marked and removed with Picard (v2.21.4). Reads mapped to the mm10 blacklist regions were removed by bedtools (v2.27.1) by intersect -v. Last, filtered reads were sieved to account for Tn5 insertion by deeptools (v3.5.3) with command alignmentSieve – ATACshift. bigWig files were generated with deeptools (v3.5.3) with command bamCoverage --binSize 10 -e --normalizeUsing CPM. For each genotype, an average binding intensity bigWig file from three replicates was generated with deeptools (v3.5.3) command bigwigAverage --binSize 10.

Peak-calling was performed with MACS2 (v2.2.7.1) on individual replicate with command macs2 callpeak --nomodel --keep-dup all. For each genotype, we generated a genotype-specific high confidence peakset by intersection of peaks called in at least two replicates. R package DiffBind (v3.0.15) and DEseq2 (v1.30.1) were then used to re-center peaks to ±200bp regions surrounding summit, generate normalized binding matrix, and differential binding matrix. We defined differentially bound peaks between each mutant and WT sample if the absolute log_2_FC was more than 2.0, corresponding to four-fold, and the FDR smaller than 1e-3.

To associate ATAC-seq peaks to genes, we used Genomic Regions Enrichment of Annotations Tool (GREAT v4.0.4) through the R package rGREAT (v1.19.2). Each peak was assigned to the closest transcription start site (TSS) within 50kb. Each gene can have one or more associated ATAC-seq defined cis-regulatory element (CRE). Overlap of ATAC-seq peaks and CRX ChIP-seq peaks were identified by pybedtools (v0.9.1) with BedTool intersect function. The compiled ATAC-seq intensity matrices can be found in the GitHub repository of this paper.

#### De novo motif searching

The mm10 fasta sequences for each genotype specific peaks were obtained using R package BSgenome (v1.66.3). *De novo* motif enrichment analysis for each set of sequences were then performed with MEME-ChIP in MEME Suite (v5.5.2) using order 1 Markov background model and default parameters. Since homeodomain motifs are relatively short and can be repetitive (e.g. K88N motif), we reported STREME found motifs, which is more sensitive than MEME to find short, repetitive motifs. In Figure 4E, PWMs for selected retinal basic helix-loop-helix (bHLH) factors were obtained directly from the JASPAR2024 website. The accession numbers are: NEUROD1 (MA1109.1), ASCL1 (MA1100.1), ATOH7 (MA1468.1). Note the ASCL1 PWM was padded at the first position for pattern alignment. All PWMs plotted can be found in Supplementary Table S4.

#### Dimeric K_50_ HD motif half-site affinity prediction

For the analysis presented in Figure S3A, we first identified dimeric K_50_ HD motifs under *Crx*^*E80A/A*^-reduced ATAC-seq peak sequences by FIMO searching (--threshold 1.0E-3) using the PWM model identified in *de novo* motif searching (section above). The identified dimeric K_50_ HD motif instances were further filtered by constraining the HD half-site core motif to match 5’-NAAN-3’. For each dimeric motif, the relative binding affinity of each 6mer half-site (Figure 1 bottom), normalized to the monomeric consensus 5’-TAATCC-3’, was calculated using a published CRX PWM model assuming position independence. The PWM models used in FIMO search and relative binding affinity calculation can be found in Supplementary Table S5.

#### HD motif type annotation for CRX bound CREs adjacent to CRX-DAGs

The list of CRX dependent-activated genes (CRX-DAGs) and their annotations were taken directly from our previous publication (17). CRX-DAG associated CRX ChIP-seq peaks were scanned for instances of K_50_ and Q_50_ HD monomeric and dimeric motifs with FIMO in MEME Suite (v5.5.2) using order 1 Markov background model and --thresh 1.0E-3. PWMs used for FIMO search can be found in Supplementary Table S5. A published CRX PWM model (27) was used to scan for K_50_ HD monomeric motifs, and a MEME found pattern under the *Crx*^*E80A/A*^-reduced ATAC-seq peaks was used to scan for K_50_ HD dimeric motifs. The RAX2 JASPAR motif MA0717.1 was used to scan for Q_50_ HD monomeric motifs, and a MEME found pattern under the *Crx*^*K88N/N*^-increased ATAC-seq peaks was used to scan for Q_50_ HD dimeric motifs. FIMO found dimeric HD motif instances were further filtered by constraining the HD half-site core motif to match 5’-NAAN-3’. To distinguish standalone monomeric motifs and high-affinity half-sites within dimeric motifs, we first searched for dimeric motif instances, masked to N, and re-run motif search for monomeric motif instances with the masked sequences. Each CRX bound CRE associated with a CRX-DAG was then annotated to have presence (1) or absence (0) of a K_50_ or Q_50_ HD motif. The full HD motif type annotation for CRX-DAG-adjacent CREs can be found in Supplementary Table S6.

#### Gene ontology analysis

Gene ontology analysis in Figures 3D and 4F were performed using R package clusterProfiler (v3.18.1) with the genome wide annotation package org.Mm.eg.db (v3.15.0). For the enrichment analysis in Figure 3D, a log_2_FC<-1 and fdr<0.05 cutoff was used to identify genes that display concordant reduction in expression and in chromatin accessibility at their nearby cis-regulatory regions. Enrichment p-values were adjusted by Benjamini-Hochberg procedure. Redundantly enriched GO terms were removed using simplify() function with parameters cutoff=0.7, by=p.adjust. The enrichment analysis results were then exported in table format and further processed for plotting with python.

#### Re-analysis of published data

The CRX ChIP-seq and bulk RNA-seq data for the WT, *Crx*^*E80A*^, *Crx*^*K88N*^, *Crx*^*R90W*^ mouse retinas from our previous publication (17) were used without alteration. To make the Figure 3E heatmap, the differential gene expression matrix was ordered by hierarchical clustering using the python package scipy (v1.11.2) with linkage method “complete” and distance metric “euclidean”.

The developmental time-course bulk ATAC-seq and RNA-seq datasets from the Aldiri et al. study (28) were obtained from GEO under accession number GSE87064. Replicates are not available for the ATAC-seq data. Otherwise, the ATAC-seq reads were processed similarly as ATAC-seq data generated in this study. The *Crx*^*E80A*^ or *Crx*^*K88N*^ consensus ATAC-seq peakset was used directly to score signal enrichment of the Aldiri ATAC-seq data. For Figures 3C, 4C, 5D-5E, accessibility z-scores were calculated using only the post-natal ages data (P0, 3, 7, 10, 14) with sklearn.preprocessing.StandardScaler from the scikit-learn package (v1.3.0). The Aldiri RNA-seq data (Figures S4A-S4B) re-processed in our previous publication (17) were used without alteration.

The VSX2 ChIP-seq data from the Bian et al. study (29) was obtained from GEO under accession number GSE196106. The reads were processed similarly as the CRX ChIP-seq data described in our previous study (17). To identify embryonic day 14.5 (E14.5) enriched, adult enriched, and shared VSX2 binding regions used in Figure 4D, a consensus peakset was first generated and subjected to non-supervised hierarchical clustering using the python package fastcluster (v1.2.6) with parameters method=single, metric=euclidean. Overlap of *Crx*^*K88N*^ differential ATAC-seq peaks and VSX2 ChIP-seq peaks were identified by pybedtools (v0.9.1) with BedTool intersect function.

#### MPRA data preprocessing

2x150bp reads from Illumina NovaSeq2000 were obtained for all samples at Genome Technology Access Center at the McDonnell Genome Institute (GTAC@MGI, WashU). Libraries prepared from un-electroporated plasmid DNAs were sequenced to 50M reads in two technical replicates (approx. 2500x). Libraries prepared from retinal *ex plant* extracted RNA (n=3/4 per genotype) and DNA (n=1 per genotype) were sequenced to an average depth of 30M reads (approx. 1600x depth). The pre-processing of MPRA reads followed the pipeline previously described (30). Briefly, the sequencing reads were demultiplexed using the inner phasing index within the P1 adapter sequences. Next, the 10bp unique CRE identifying barcodes were extracted and counted with customized python scripts. The compiled MPRA CRE barcode count matrix and annotation table can be found at the GitHub repository for this paper.

#### MPRA regulatory activity calculation

By design, each testing CRE was represented by four unique barcodes (Figure S6B). One or more of these barcoded CREs can dropout in the process of cloning, or due to low activity. To ensure quality measurement, we dropped 1) barcodes with less than 50 reads in both plasmid libraries; 2) CREs represented by less than 3 unique barcodes after step 1). Then, we normalized the reads from the plasmid libraries by counts-per-million paradigm without adding pseudocount. Reads from the retinal *ex plant* extracted RNA and DNA libraries were normalized similarly by adding 1 pseudocount to each barcode. Counts of individual RNA libraries were normalized by the average counts of the plasmid libraries to obtain “raw activity score” for individual barcode. Average raw activity scores by unique barcodes for individual genotype was calculated by averaging the genotype replicates. Then, average raw activity scores by unique testing CREs for individual genotype was calculated by averaging the barcode raw activity scores for each CRE. A coefficient of variation threshold of 1.0 was used to filter out CREs whose barcode activity vary greatly among genotype replicates. Last, to calculate “regulatory activity score”, the raw activity score of each CRE was normalized to the average raw activity score of all scrambled control CREs.

#### MPRA HD motif activity analysis

In the MPRA library design, each testing genomic CRE sequence was scanned for the presence of monomeric and dimeric K_50_ and Q_50_ HD motifs following the same pipeline as HD motif type annotation for CRX bound CREs (section above). A less stringent --thresh 2.5E-3 for FIMO search was used to capture more non-consensus motifs. Both FIMO found monomeric and dimeric HD motif instances were further filtered by constraining the HD monomeric site or half-site core motif to match 5’-TAA-3’ at the first three positions. If a testing genomic CRE sequence (WT) contains HD motif(s), one or more mutated CRE versions were generated: 1) mutM: all and only the monomeric HD motifs mutated; 2) mutD: all and only the dimeric HD motifs mutated; 3) mutDM: all monomeric and dimeric HD motifs mutated if both are present. To minimize perturbations of nearby/overlapping TF motifs, a single substitution 5’-TAA-3’ to 5’-TAC-3’ was introduced to disrupt HD motifs. This substitution has been found in EMSA to abolish CRX HD binding to DNA (31). K_50_ or Q_50_ type HD motifs were not differentiated due to the complexity of library design and the differential binding of CRX WT vs K88N proteins on these two types of motifs. The “HD motif activity score” (Figures 7B-7C) was calculated as the “regulatory activity score” difference between of a mutant CRE version and its matched genomic CRE sequence. The HD motif position information and compiled HD motif activity score matrix can be found at the GitHub repository for this paper.

#### Statistical analysis

One-way ANOVA with Turkey honestly significant difference (HSD) tests in Figures 1E and S1D were performed with python packages scipy (v1.11.2) and scikit_posthocs (v0.7.0). Fisher’s exact test in Figure 4D was performed with python package scipy (v1.11.2). Mann-Whitney-Wilcoxon tests in Figures 7B-7C were performed with scipy through python package statannotations (v0.4.4).

## Supplementary data

Supplementary Figures S1-S6 Supplementary Tables S1-S6

## Author contributions

S. Chen and Y. Zheng conceived the study. S. Chen and G.D. Stormo supervised the study. S. Chen and Y. Zheng designed the experiments. Y. Zheng performed all the experiments, data analysis and visualization. G.D. Stormo assisted in Coop-seq data analysis. Y. Zheng wrote the original draft. S. Chen, G.D. Stormo, and Y. Zheng revised the manuscript. All authors read and approved the final manuscript.

## Acknowledgment

We thank Mingyan Yang for technical assistance; Mike Casey from the Molecular Genetics Service Core for generating luciferase reporter assay plasmids and MPRA oligo library; J. Hoisington-Lopez and M. Crosby from DNA Sequencing Innovation Lab at the Center for Genome Sciences & Systems Biology (CGS&SB) as well as Genome Technology Access Center at the McDonnell Genome Institute (GTAC@MGI) for sequencing assistance. We also thank Mr. Artur Widlak for the generous gift from Widłak Family CRX Research Fund.

## KEY RESOURCES TABLE

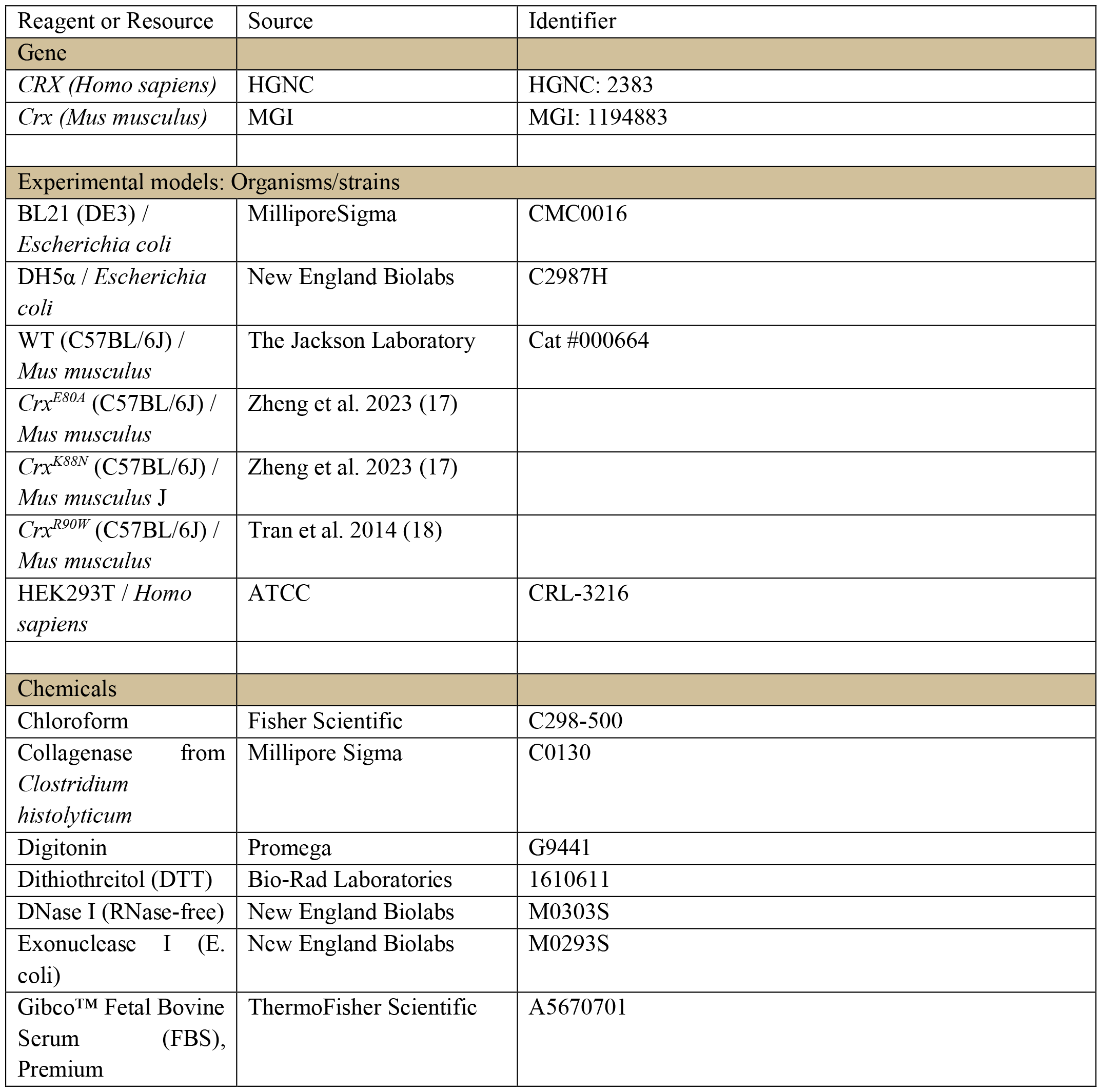
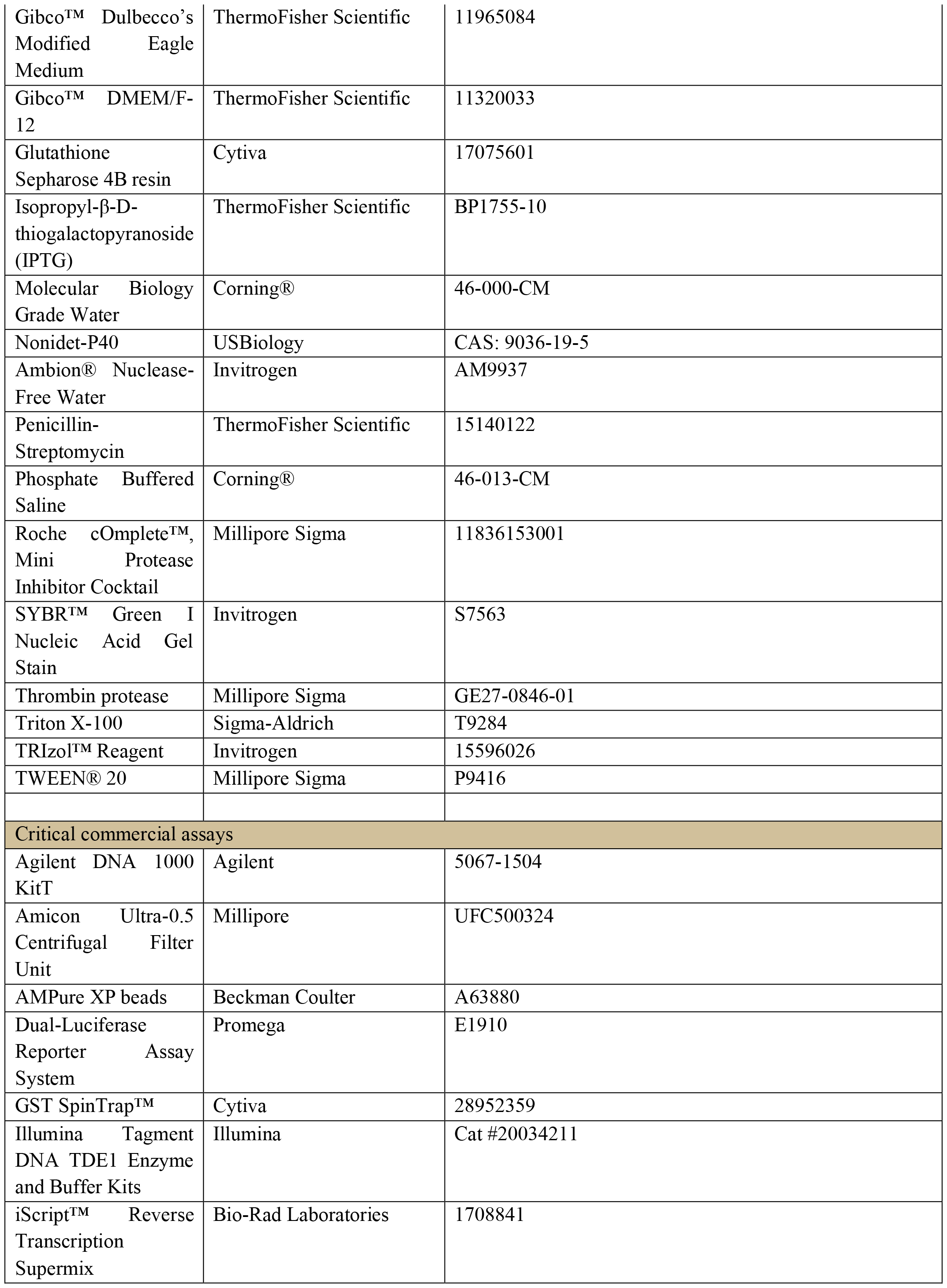
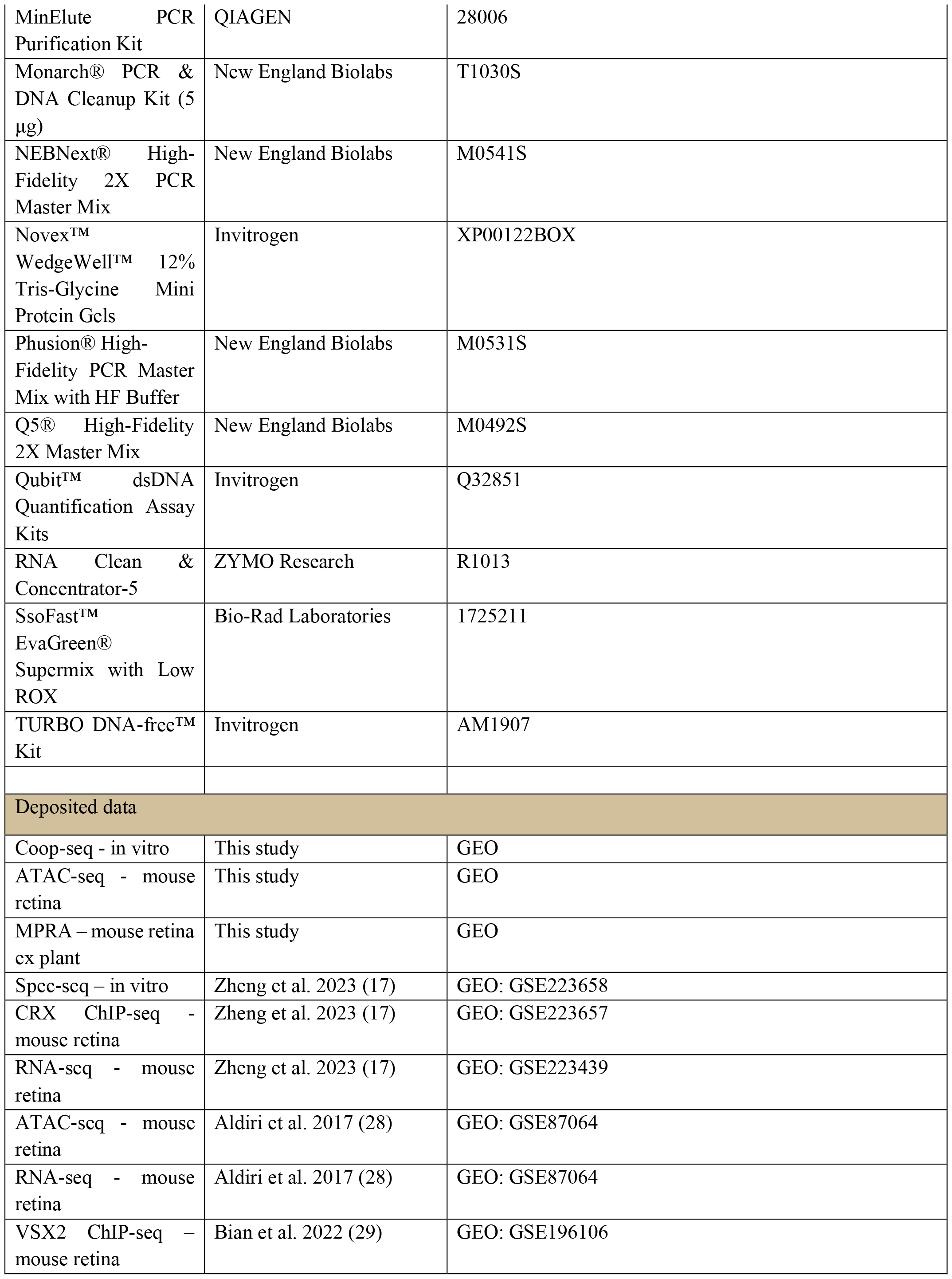
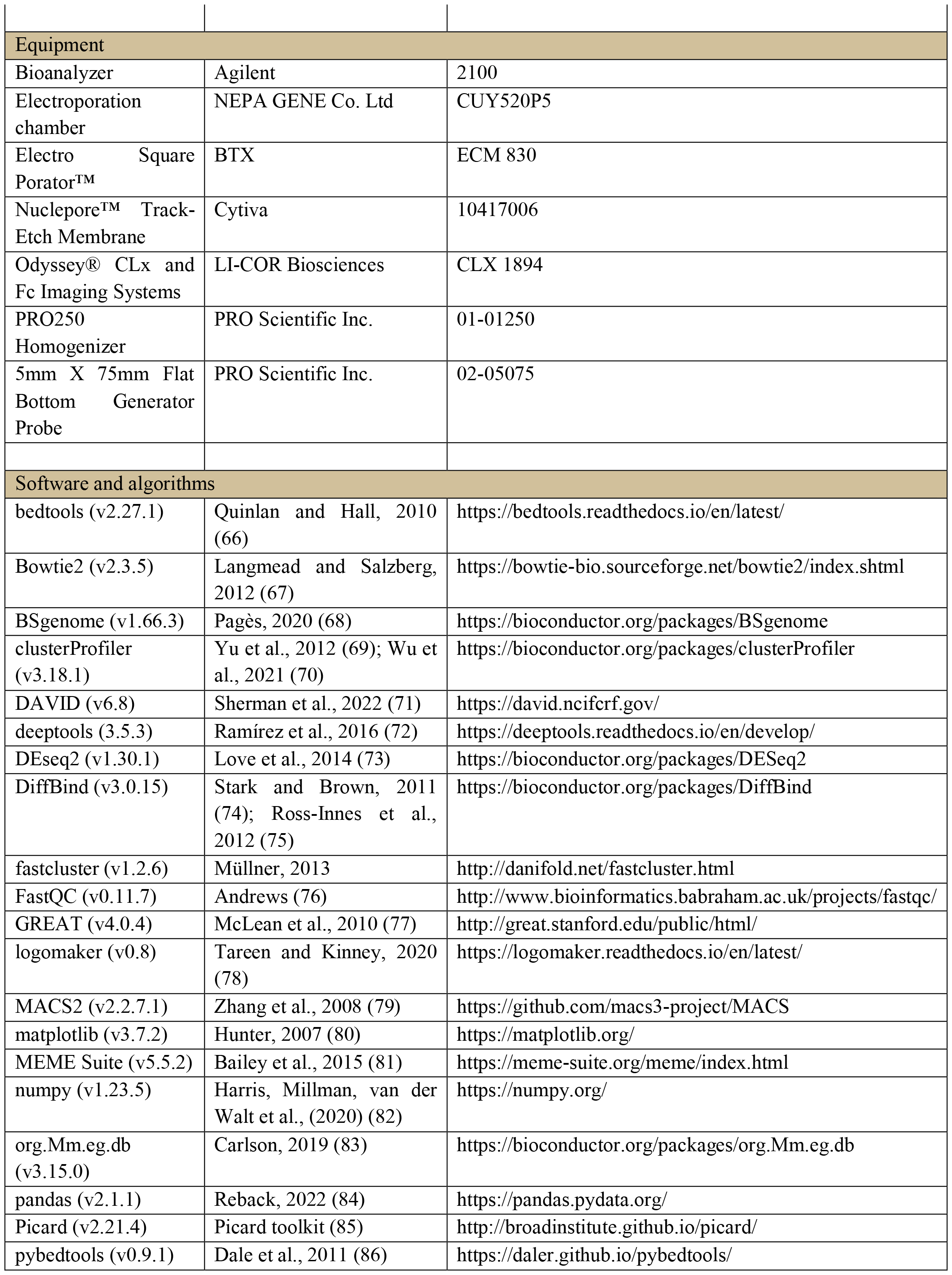
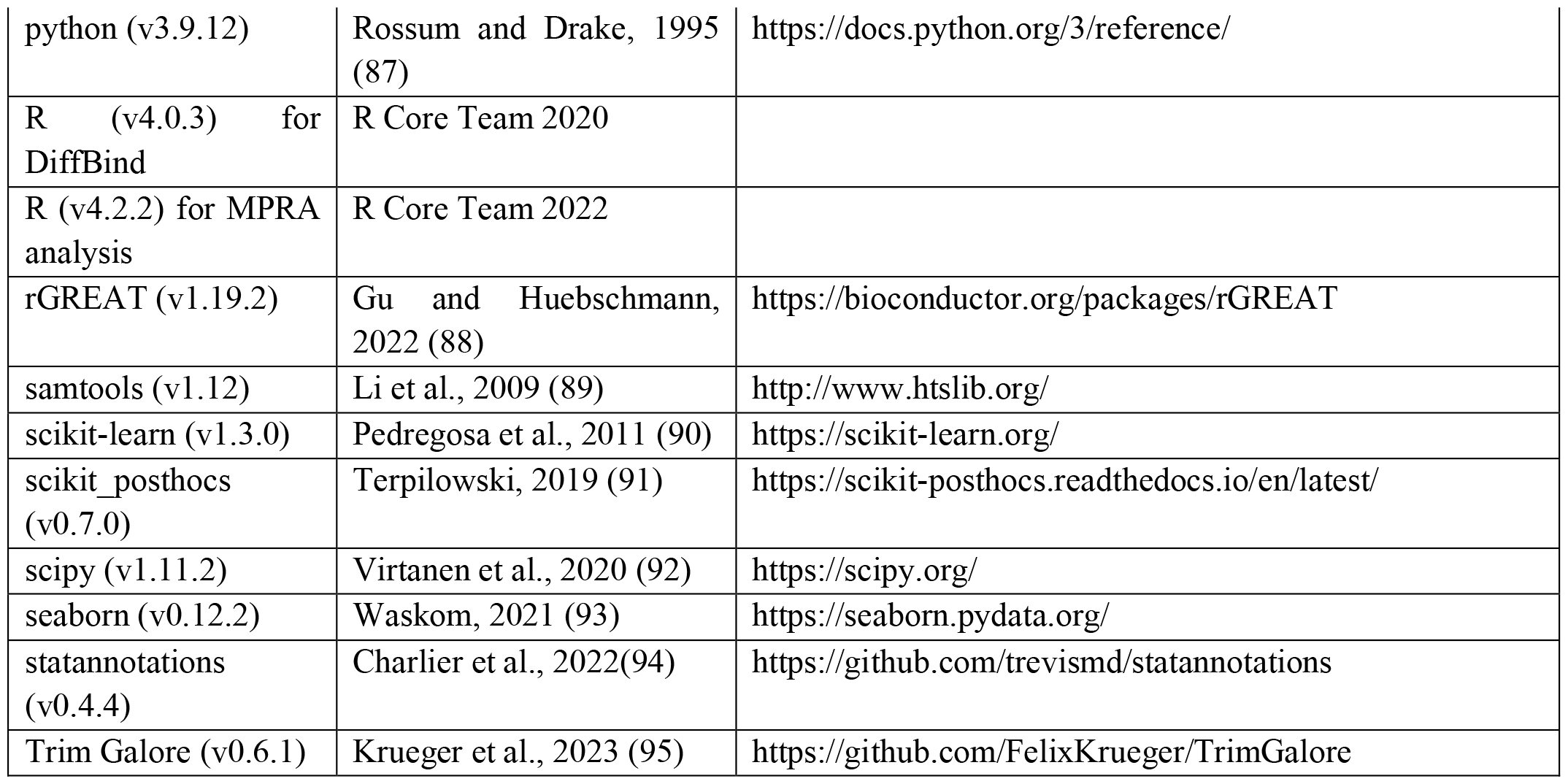

---

**Figure S1.**
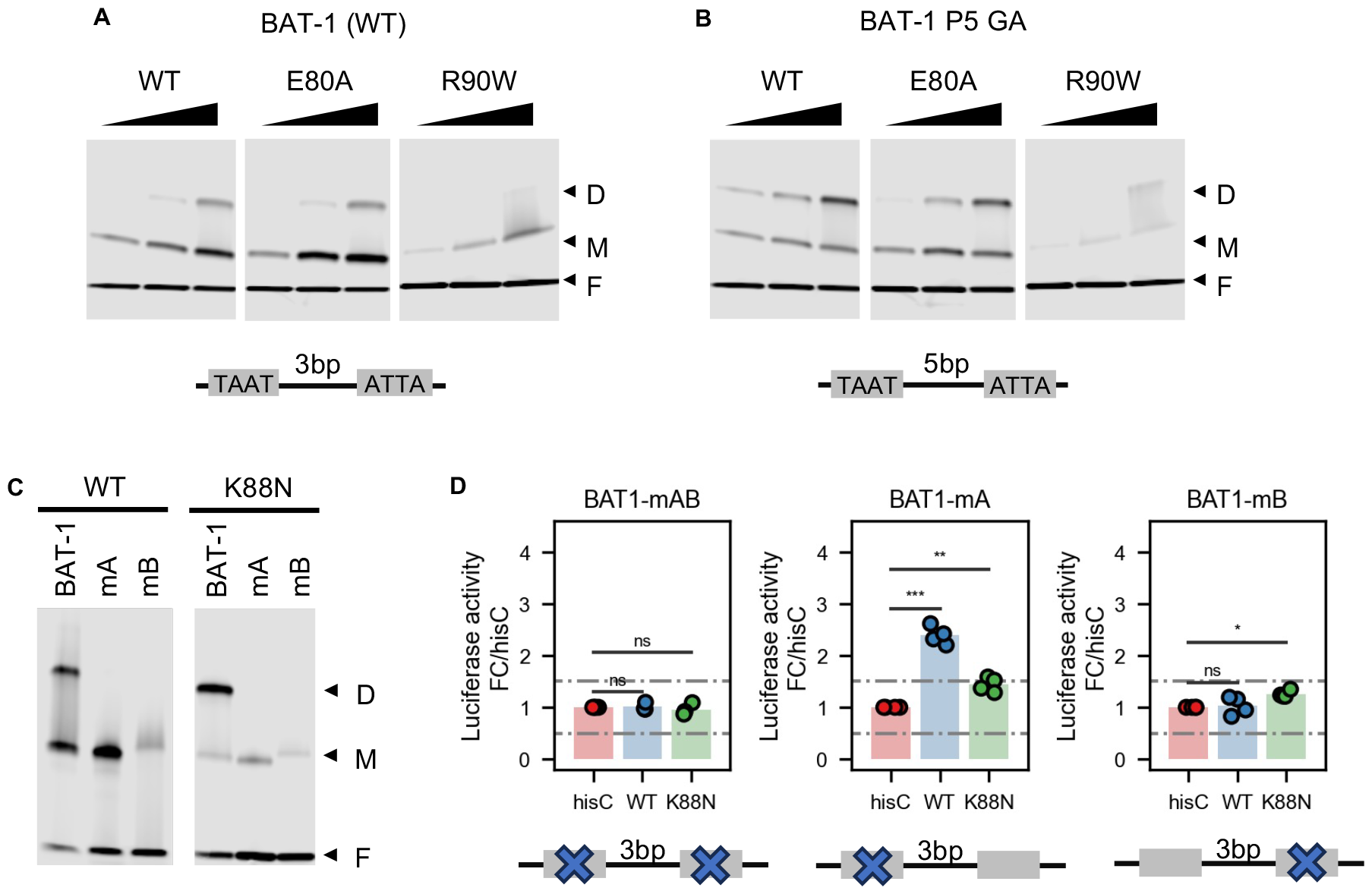
P3 configuration and intact half-sites are required for K88N HD cooperative binding and transactivation activity. **A.&B**. EMSA gel images showing increasing amount of WT, E80A, R90W HD peptides bound to a fixed amount of BAT-1 (WT) or P5 probes. The cartoon underneath each gel image shows the dimeric HD motif configuration and is labeled with the spacer length. Note the WT HD EMSA gel images are the same as in Figures 1C-1D. **C**. EMSA gel images showing WT and K88N HD peptides bound to BAT-1 (WT), mA and mB probes. **D**. Barcharts of luciferase reporter assays comparing CRX WT and K88N transactivation activity at BAT-1 variant enhancer sequences. The cartoon underneath each barchart shows the HD core motif mutated. *p-values* of one-way ANOVA are annotated. ns: >5e-2; *: <=5e-2; **: <=1e-2; ***: <=1e-3.

**Figure S2.**
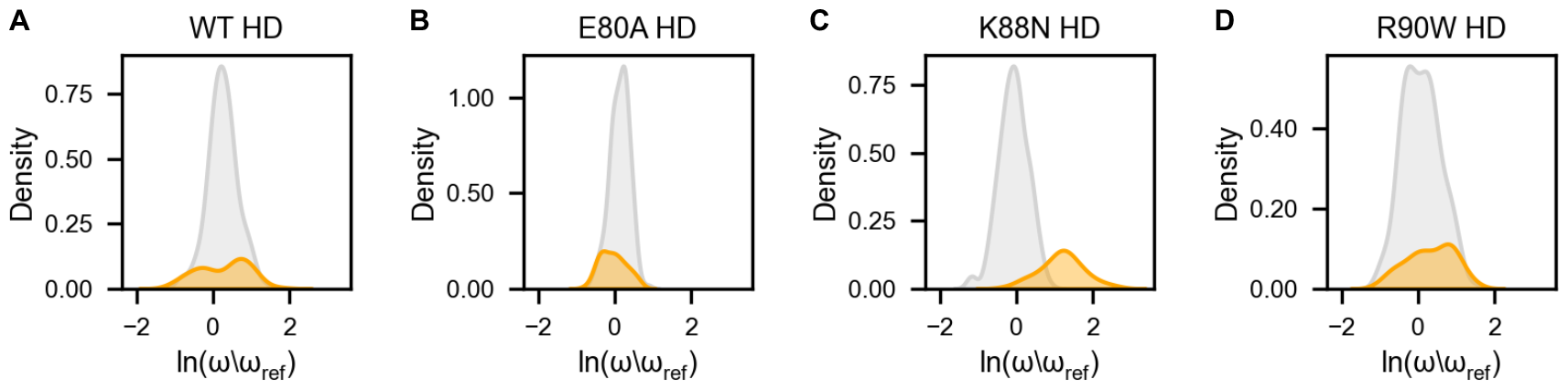
WT and mutant CRX HDs demonstrate similar cooperativity profiles at P5 library but distinct cooperativity profiles at P3 library. **A.-D**. Histograms depicting the distribution of CRX HD relative cooperativity on members of the P5 library (grey) and the P3 library (orange). Relative cooperativity against the reference sequence TAATGCGCTATTA is plotted (ω/ω_ref_). Note the relative cooperativity is presented in the Logarithmic scale. The full relative cooperativity matrix can be found in Supplementary Table S2.

**Figure S3.**
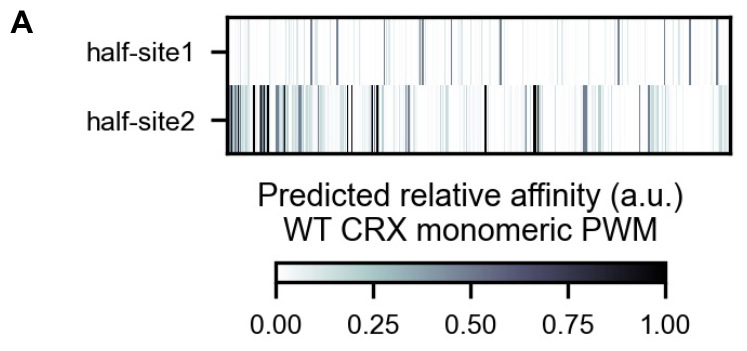
Dimeric K_50_ HD motifs in *Crx*^*E80A/A*^-reduced ATAC-seq peaks composed of low-affinity half-sites. **A**. Heatmap depicting the predicted half-site affinity of FIMO-found dimeric K_50_ HD motifs under *Crx*^*E80A/A*^-reduced ATAC-seq peaks. The dimeric motifs are ordered by FIMO p-values. The half-site relative affinity, normalized to consensus TAATCC, is predicted using the published WT CRX PWM model (Supplementary Table S4, Methods).

**Figure S4.**
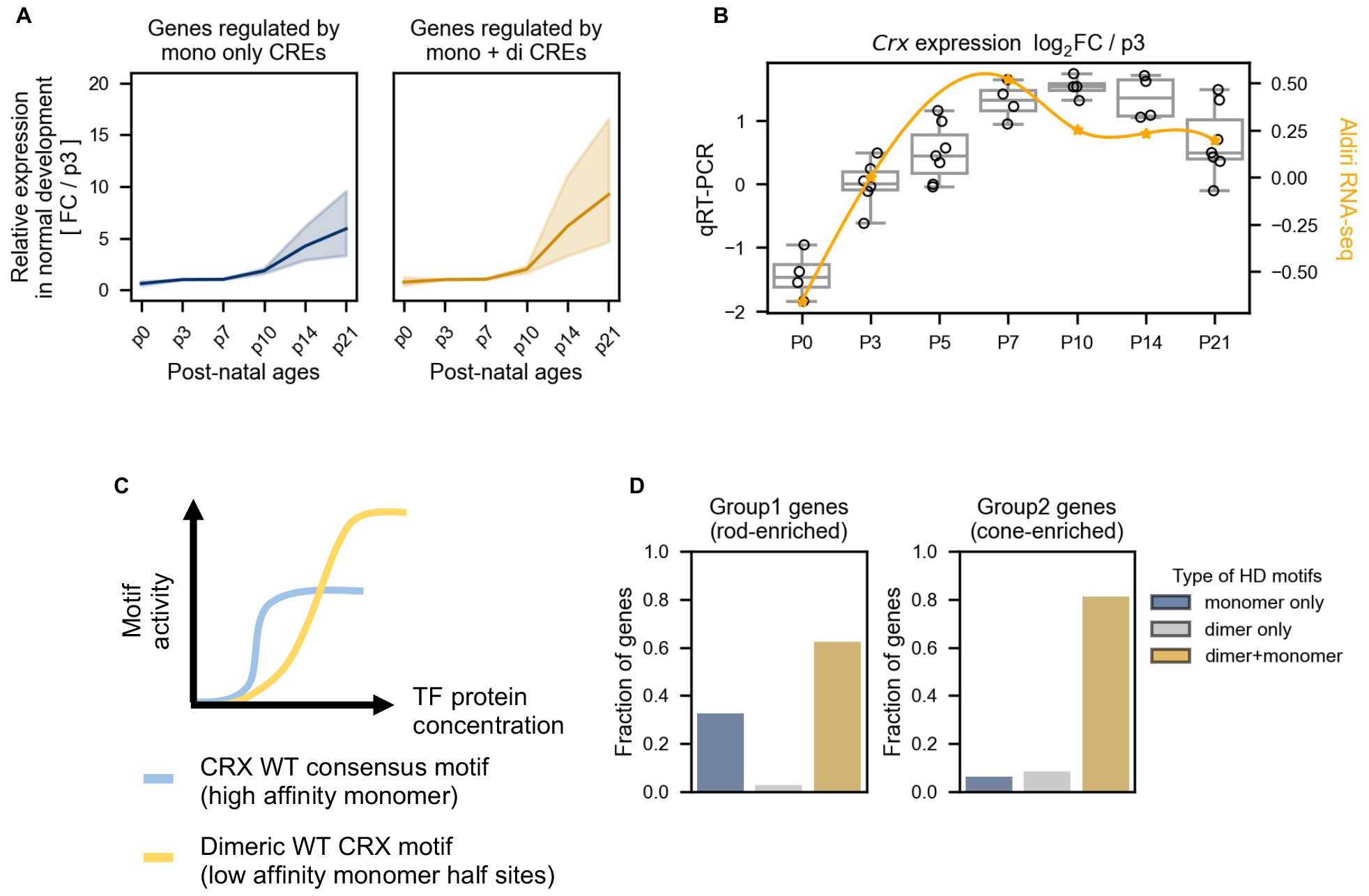
Dimeric K_50_ HD motifs are essential for late-stage photoreceptor gene expression. **A**. Line plots showing the expression dynamics of genes associated with CREs with only monomeric or both monomeric and dimeric K_50_ HD motifs. Expression levels are normalized to P3 and represented by fold-change. **B**. Dual-axes box and line plot showing the *Crx* mRNA developmental expression patterns measured by qRT-PCR (left axis, box and strip plot) and by whole-retina RNA-seq (right axis, line and scatter plot). Expression levels are normalized to post-natal day 3 (P3) and presented by log-fold-change. The developmental RNA-seq data in **S4A-S4B** is from Aldiri et al. 2017. **C**. Diagram depicting the stage-specific activity model of CRX WT high-affinity monomeric motifs and dimeric motifs that are composed of low-affinity half-sites. **D**. Barcharts showing the distribution of cone-vs rod-enriched CRX-DAGs associated with CREs containing monomeric, dimeric, or both types of K_50_ HD motifs. The definition of Group1 and 2 genes is taken directly from our previous publication Zheng et al. 2023.

**Figure S5.**
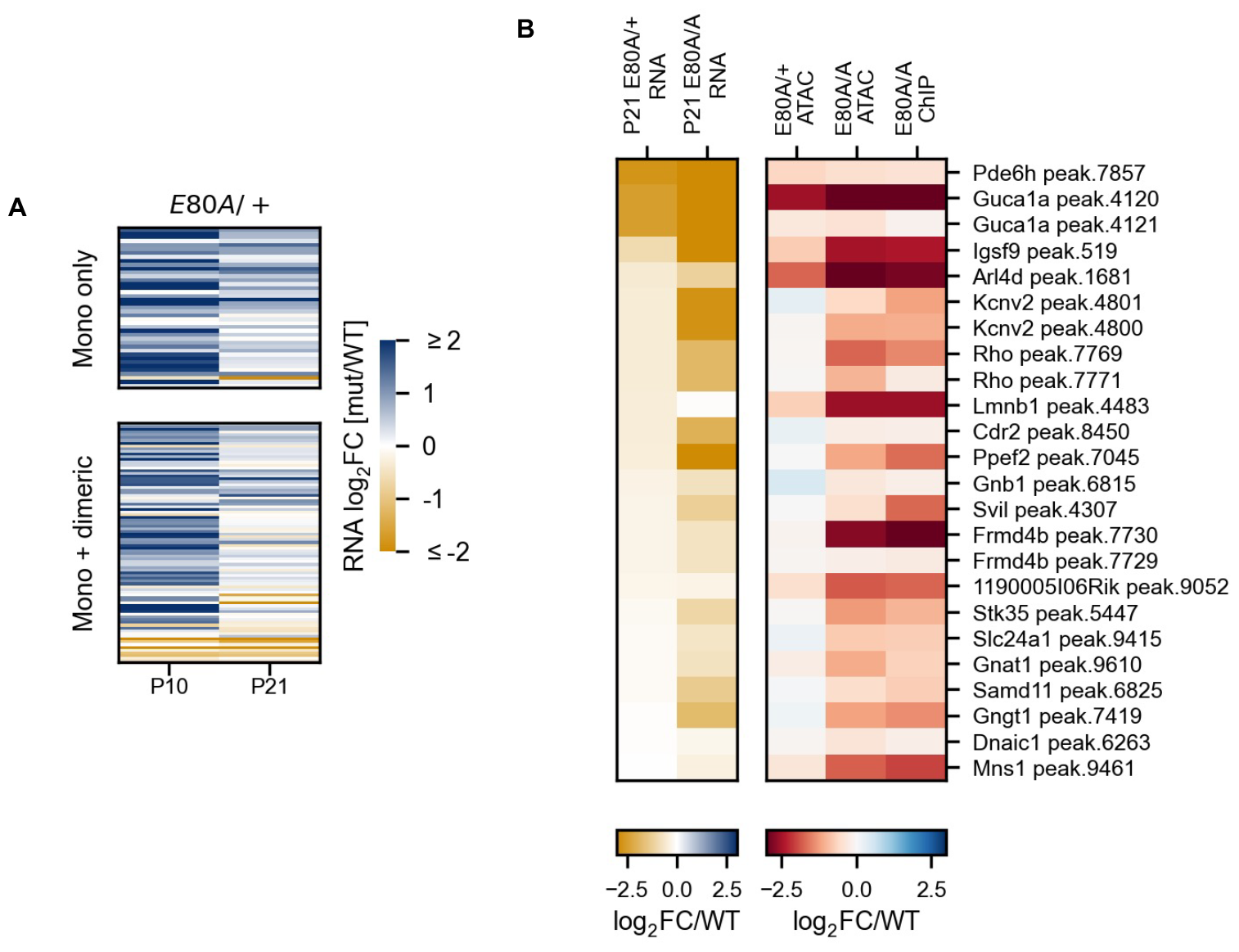
CRX-DAGs regulated by dimeric K_50_ HD motifs are selectively down-regulated in *Crx*^*E80A*^ mutant retinas. **A**. Heatmap comparing the CRX-DAG expression changes in the *Crx*^*E80A/+*^ retinas at ages of post-natal day 10 (P10) and day 21 (P21). The gene sets in the heatmaps are as defined in **Figure 6A**. The order of the genes is identical to that in **Figure 6B. B**. Heatmaps showing down-regulated genes with correspondingly decreased chromatin accessibility at associated CREs in the *Crx*^*E80A*^ retinas. The genes and CREs are ordered first by RNA-seq logFC than by ATAC-seq logFC in the *Crx*^*E80A/A*^ retinas. The peak.id denotes the unique CRX ChIP-seq peak associated with a gene. The full CRX bound CRE-gene association matrix can be found at Supplementary Table S6.

**Figure S6.**
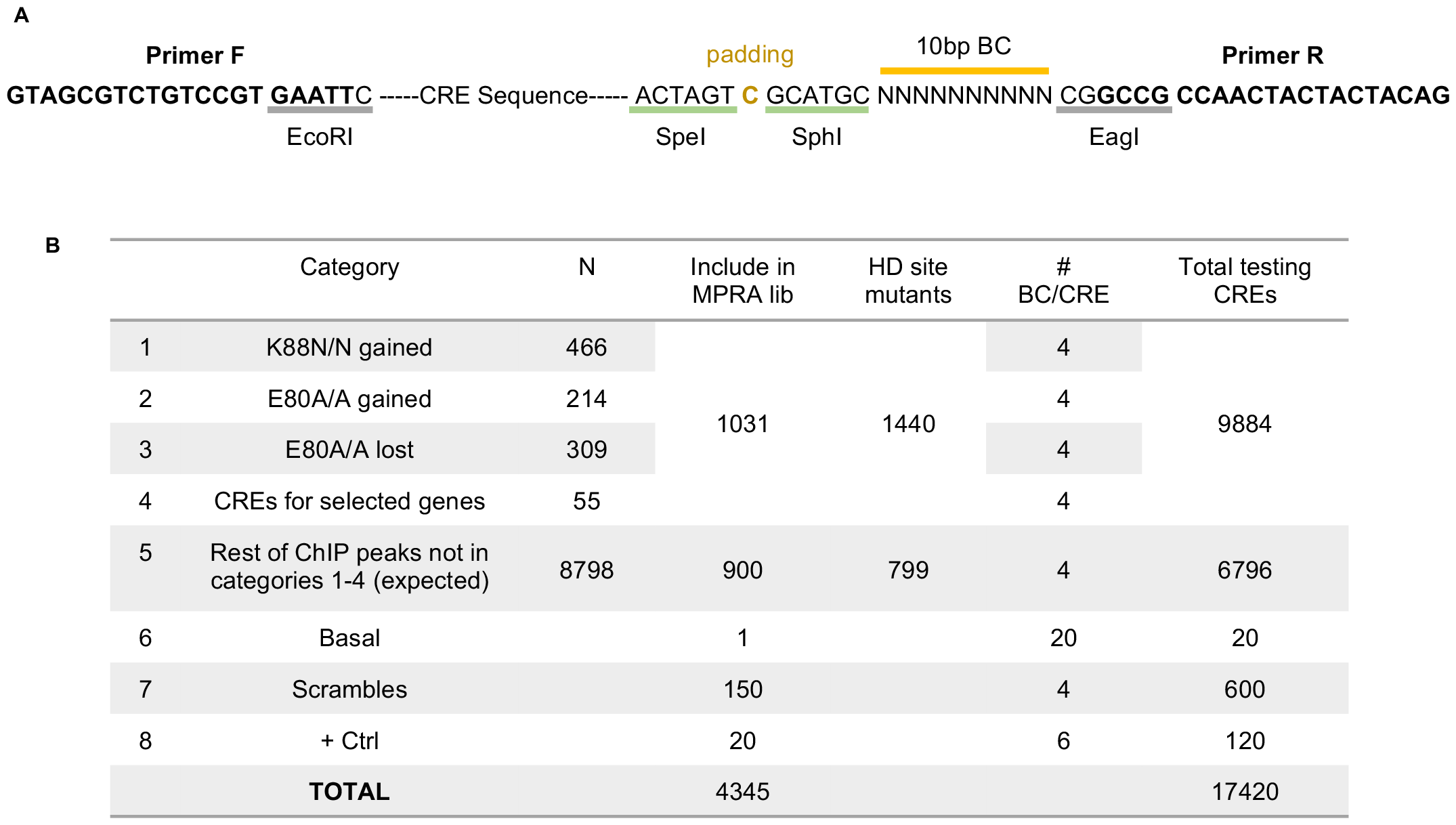
MPRA oligo design and library composition. **A**. Diagram of a typical testing MPRA oligo. The restriction enzyme cut sites used for cloning, primer sites for amplification, and testing CRE and barcode locations are noted. **B**. Table displaying the composition of MPRA library designed and ordered. The gained and lost categories were defined by comparing the ChIP-seq signal intensity (Methods, MPRA GitHub repository).

